# Structured Pooling Improves Detection of Rare Regulatory Mutations in Population-Scale Reporter Assays

**DOI:** 10.64898/2026.03.27.714794

**Authors:** Katherine Dura, Keith Siklenka, Kari Strause, Shauna Morrow, Chuangchuang Zhang, Alejandro Barrera, Andrew S. Allen, Timothy E. Reddy, William H. Majoros

## Abstract

Identifying genetic variants in noncoding DNA that impact gene expression and thereby contribute to disease risk remains a difficult but important challenge in genomic medicine. Modern reporter assays such as STARR-seq and MPRA provide an efficient and effective means of testing, in very high throughput, millions of variants captured directly from patient genomes. While these assays have previously been scaled to whole genomes and, separately, to populations, we report findings from the first whole-genome population-scale STARR-seq experiment performed on 100 individuals. In order to achieve that scale we devised a novel experimental design that partitions samples into pools so as to increase allele frequencies within pools and thereby reduce expected dropout and increase signal-to-noise ratio in experimental readouts. We show that this design produces more accurate estimates of variant effect sizes, and we provide a Bayesian model for robust estimation of those effect sizes that also reports full posterior distributions for assessment of confidence in estimates. Together, these methodological innovations facilitate the detection of functional regulatory variants, particularly rare variants, with much higher accuracy and at greater scale than previously possible. We demonstrate the utility of this approach on the task of functional annotation of quantitative trait loci such as eQTLs and caQTLs, and show concordance with patterns of constraint in transcription factor binding profiles.

## 1. Introduction

A major challenge in functional genomics is the systematic identification of genetic variants that contribute causally to disease risk. In pursuit of that goal, a common approach is to systematically search for correlations between clinical traits and the alleles or haplotypes present in a cohort within each region across the entire genome. A major finding of such genome-wide association studies (GWAS) that have been carried out to date is that sizeable portions of their signals map to the noncoding portion of the genome, potentially indicating a large role for genetic variation in the cis-regulatory machinery in explaining variation in human phenotypes^1–8^. Indeed, a number of examples exist of noncoding regulatory variants that have been demonstrated to mechanistically interfere with transcription initiation, thereby potentially contributing to disease^9–13^.

While GWAS studies can be very effective at identifying haplotypes correlated with traits, their results can be challenging to interpret when detected signals are located in non-coding regions of the genome^14^. Because of the phenomenon of linkage disequilibrium (LD), the regions implicated in association signals can contain numerous variants all correlated with a trait, but which likely do not all contribute causally to phenotypic outcomes. Thus, resolution is often insufficient in these studies to narrow the set of candidate variants to a manageable set for experimental validation. Moreover, the issue of resolution can be particularly limiting for noncoding signals, as cis regulatory elements and the protein binding sites within them are generally much smaller than typical protein-coding genes, making the task of mechanistic interpretation of noncoding association signals very challenging.

To address this problem, various assays have been developed in an attempt to more finely dissect contributions of genetic variation to dysfunction in regulatory mechanisms^12,13,15,16^. In massively parallel reporter assays^17^ and related experiments such as STARR-seq^18^, a library of DNA plasmids is generated and transfected into cells to produce an associated library of mRNA transcripts produced from a common reporter gene. These DNA and RNA libraries are both sequenced, and the alleles represented among the sequencing reads are counted. By comparing the corresponding allele counts in both libraries for each assayed element, one is able to compare the relative transcriptional activity resulting from the different alleles for a single variant (Fig. 1a).

**Figure 1:**
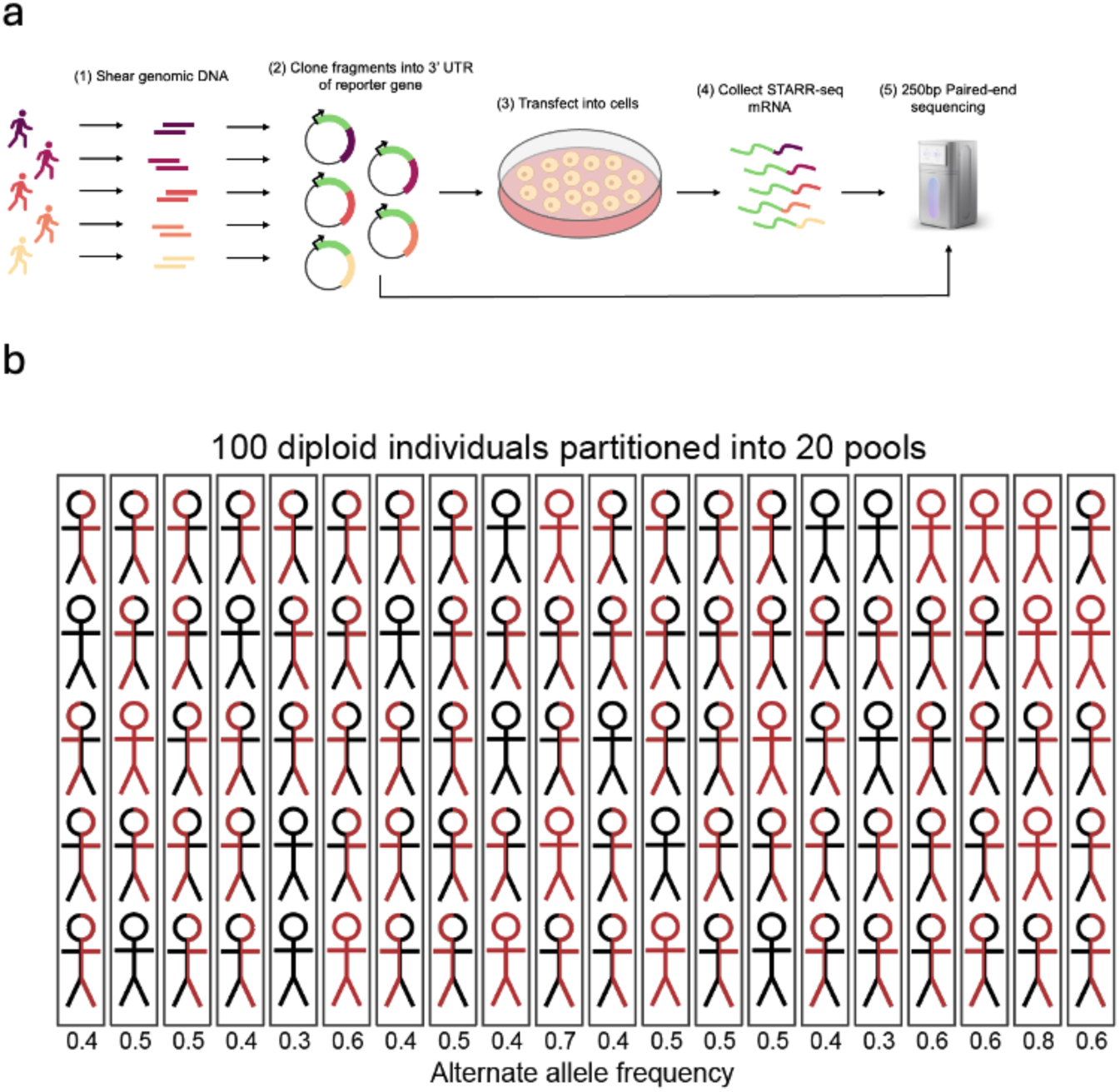
(a) This panel was adapted from Dura et. al. 2025. Schematic of the STARR-seq experiment. Genomic fragments are sheared from samples and cloned into the 3’UTR of a reporter gene. Those fragments are sequenced to determine the allele frequency of each variant in the DNA library. Fragments are also transfected into cells, where STARR-seq mRNA is produced. Those mRNA are sequenced to determine allele frequencies in the RNA library for comparison to their frequencies in DNA and statistical inference of regulatory effects. (b) Structured pooling replaces the traditional single-pool design with multiple pools containing disjoint subsets of samples, resulting in measurements of allele frequency in each pool for both DNA and RNA.

Importantly, these assays can be performed at the whole-genome scale^19^, so that functional annotation of regulatory variants can be achieved systematically across the entire regulatory genome in a single experiment. To the extent that the DNA inserts in the plasmids are often smaller than LD blocks, measurements from these experiments can more finely annotate variant contributions to regulatory signal and thus potentially to trait associations, because those inserts are tested out-of-genome via episomal constructs, thus eliminating confounding factors in the local genomic context. In addition, genomic material from multiple individuals can be pooled into a single library, to drastically increase the amount of genetic variation that is simultaneously assayed^20,21^.

A major limitation of that approach, when genomic DNA from multiple samples is pooled into a single library, is that the relative frequency of rare alleles present in the library can dramatically decrease as the cohort size is increased. For example, personal variants will have an expected relative frequency of 1/(2N) in a plasmid library formed from N diploid individuals. Assuming a fixed library size and constant sequencing effort, each variant thus experiences an increased risk of undergoing dropout and therefore being absent from the final experimental readout as the assayed population grows--i.e., as N increases. Thus, while pooling larger numbers of individuals in a single experiment will generally result in larger numbers of variants being represented in the input samples, it incurs a concomitant risk of losing more variants due to dropout during library construction, if the plasmid library size is kept fixed. While this could in principle be counteracted by a corresponding increase in library size--e.g., scaling by a factor of N by constructing a plasmid library for every individual in the cohort--that approach becomes prohibitively costly for large populations.

Here we introduce a new experimental design for whole-genome population-scale reporter assays that addresses this problem, together with a statistical model designed to produce more accurate estimates of variant effects from these experiments. The key innovation is to partition samples into disjoint pools, each of which is used to construct a separate plasmid library for independent transfection (Fig. 1b). Partitioning in this way naturally elevates the effective allele frequencies of rare variants within each pool, thus mitigating the incidence of dropout. For a cohort of 100 diploid individuals partitioned into 20 pools, a personal variant with frequency 1/200 in the cohort will have a frequency of 1/10 within the pool that receives that sample. However, because of drift in allele frequencies that occurs due to stochasticity in plasmid formation and transfection, actual frequencies can vary during the experiment. We thus derive a new statistical model that estimates actual frequencies across pools and uses these estimates in a Bayesian manner to produce full posterior distributions of effect size that provide information about the variances in those estimates, and thus allow one to gauge the confidence in estimated variant effects.

Applying this experimental design to 100 samples from the Thousand Genomes Project, we were able to assay ∼16.9 million variants genome-wide (∼1.5 million in high-activity STARR-seq peaks) for regulatory effect. Using data from this experiment we show that estimates of variant effects are highly concordant with both chromatin quantitative trait loci (caQTLs) and expression QTLs (eQTLS) from the African Functional Genomics Resource^22^, as well as with patterns of constraint at individual positions within transcription factor (TF) binding profiles. We also demonstrate the use of these functional annotations to dissect association signals in individual regions from eQTL and caQTL studies.

## 2. Results

### 2.1 Structured pooling enhances heterogeneity and improves accuracy of effect estimates

In previous work we showed that increasing the number of replicates in a STARR-seq experiment can result in larger heterogeneity (i.e., variance) in allele frequency between replicates, which, due to properties of the Poisson-binomial distribution, results in reduced variance in allelic read counts^23^. We also demonstrated empirically a trend toward lower estimation error as result of that increased heterogeneity. In the present study, we explore the potential for further increasing the heterogeneity (and estimation accuracy) by using disjoint sample pools instead of replicates--i.e., where each assayed pool is formed from different samples and thus may have different genotype frequencies.

Consistent with our previous results, we observed in simulation studies an overall trend toward increasing heterogeneity in allele frequency when increasing the number of simulated replicates (Fig. 2a; Suppl. Fig. 2). We also saw a strong trend toward increasing heterogeneity in multi-pool experiments when increasing the number of simulated pools. Importantly, the increase was substantially larger for the pooled structure (Fig. 2b). As expected, these trends produced lower variance in alternate allele read counts (Fig. 2c), lower variance in alternate allele frequency (Fig. 2d), lower variance in estimates of effect size θ (Fig. 2e), and lower error in estimates of effect size (Fig. 2f) for the naive estimator computed via a simple ratio of read counts, without any additional statistical modeling.

**Figure 2.**
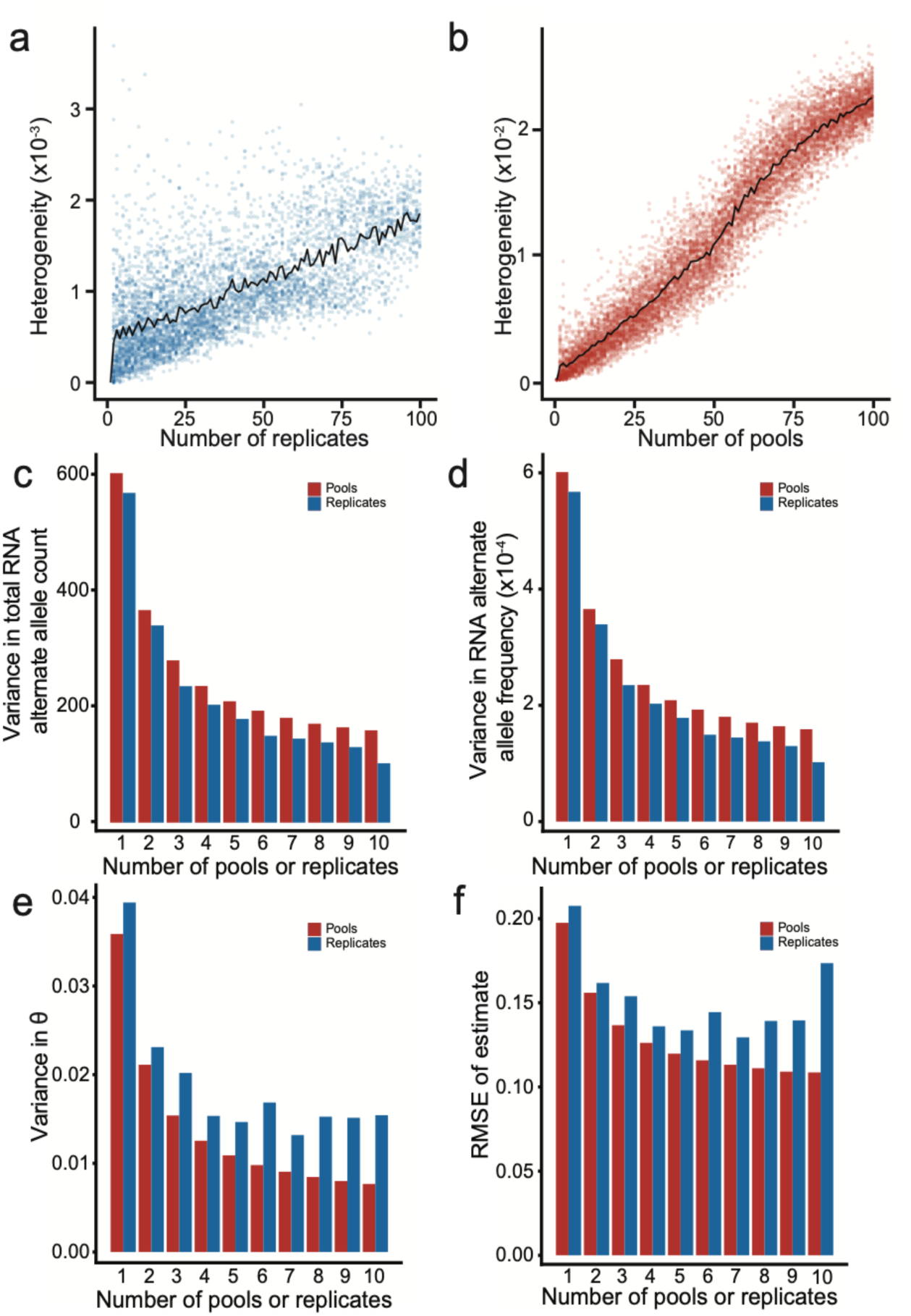
(a-b) Heterogeneity is represented as a function of the number of (a) replicates or (b) pools of a single experiment. In this case, there are 100 total samples assayed across all replicates or pools. The number of replicates or pools is represented on the X-axis. Heterogeneity in RNA is defined as the sample variance in the alternate allele frequency in the RNA, represented on the Y-axis. The black trend line indicates the average heterogeneity at each number of replicates assayed. (c-f) Comparisons of the effect of heterogeneity on estimation of effects when dividing samples among multiple pools (red) or multiple replicates (blue). Samples are divided up to 10 times. The X-axis represents the number of pools represented by the simulation. The Y-axis represents the statistic being compared, including (c) the variance in the alternate allele count in the RNA, (d) the variance in the alternate allele frequency in the RNA, (e) the variance in the effect size estimate, and (f) the RMSE Of the effect size estimate.

Indeed, in our simulations the heterogeneity of the allele frequency in RNA when increasing the number of distinct pools ranged up to 3e-2, a ∼10-fold increase in heterogeneity relative to increasing the number of replicates (Fig. 2b). Based on Wilson’s estimator of binomial variance between replicates in the experimental data for the first pool, the true allele frequency fell outside the 95% confidence interval in 25.3% of RNA replicates (Suppl. Fig. 1a; N=11,258,674), versus 8.4% in DNA replicates (Suppl. Fig. 1b; N=17,029,310), indicating that allele frequency likely drifts somewhat during library preparation, but that it drifts to a much greater degree during or after transfection (due presumably to a combination of biological variability and individual variant effects), producing substantial heterogeneity in allele frequency in RNA counts.

### 2.2 Bayesian estimation further benefits from structured pooling

We further tested the effect of multi-pool experimental strategies by applying our updated Bayesian model, BIRDbath, to data simulated to have between 1 and 20 pools, over a range of allele frequencies. A potential benefit of the structured pooling design is that the minor allele frequency of variants in one pool can be inflated relative to the entire sample cohort. For example, for a cohort of 100 diploids a personal variant would have a minor allele frequency of 0.005 among the entire cohort; partitioning into 20 pools of 5 individuals would increase this frequency to 0.1 in a single pool containing the variant (Fig. 1b). For this reason, we hypothesized that partitioning the dataset into disjoint pools may provide for better effect predictions for rare variants.

Consistent with that hypothesis, we observed that simulating 20 pools of 5 individuals resulted in more accurate estimates of effect size, measured by MSE and Spearman rank correlation, as compared to applying the model to collapsed data in which allelic read counts were summed across pools and presented to the original BIRD model as counts from a single pool with a single replicate (Fig. 3a,b,c; Suppl. Fig. 3,4,5). MSE differences were between 0.19 and 0.3, indicating a large reduction in estimation error. The advantage due to the multi-pool approach increased as allele frequency decreased, with common variants seeing an improvement in correlation of ∼0.2 and rare variants seeing an improvement of ∼0.3. These results suggest that structured pooling can be beneficial even for the estimation of rare variant effects. Improvements in estimation accuracy were due to decreases in both false positives and false negatives (Suppl. Fig. 5). We further investigated the effect of imposing different numbers of pools while keeping the total number of individuals and per-variant sequencing depth constant (Fig 3d,e, Suppl. Fig. 4). While MSE continues to decrease as the number of pools is increased past 5, we posit that in practice those additional decreases will in most cases provide only marginal difference in utility for annotation of functional variants.

**Figure 3.**
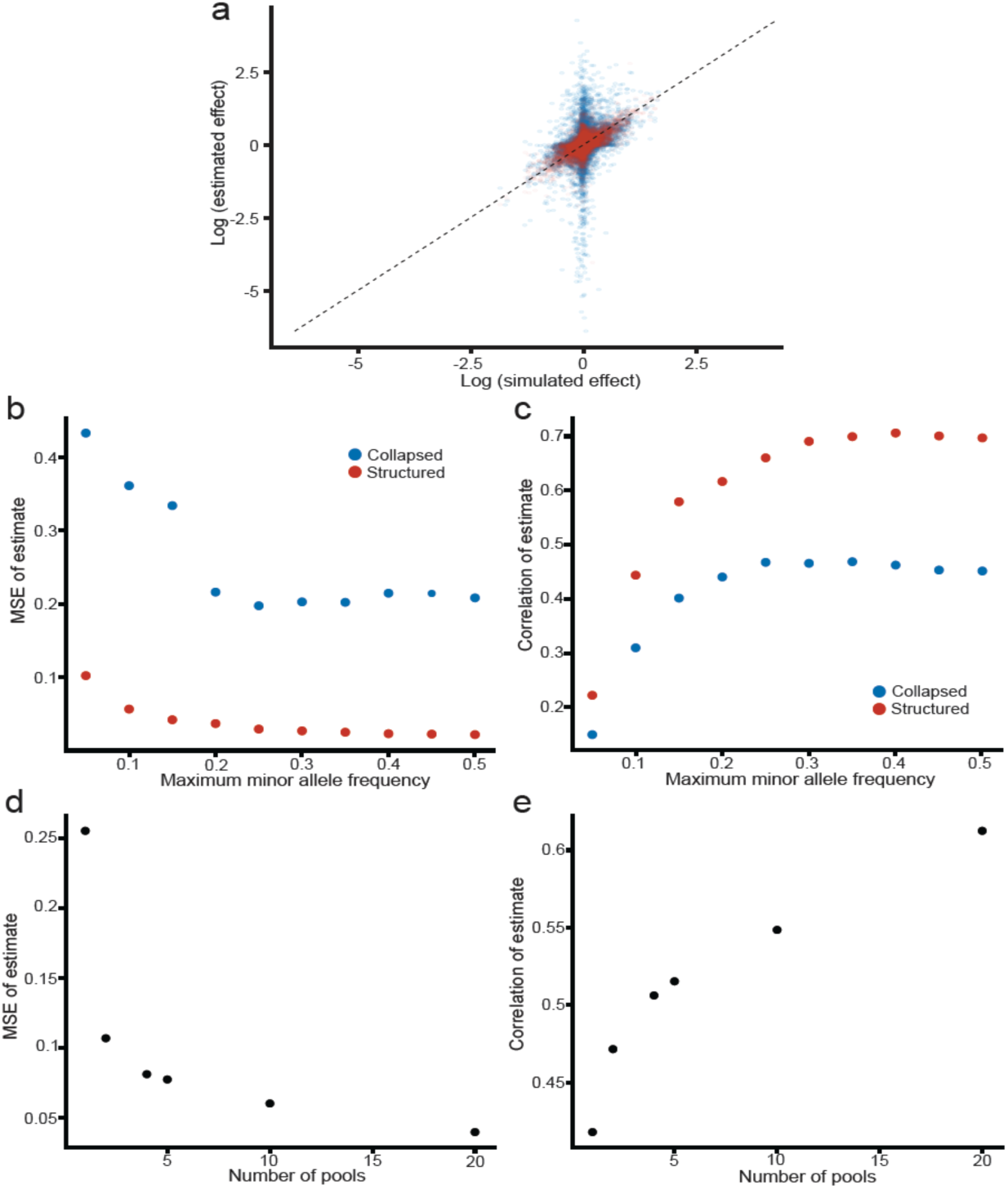
(a) Correlation between simulated effect sizes and effect sizes estimated using the BIRDbath model and the BIRD model, represented in blue and red respectively. The x-axis represents the log-transformed simulated effect sizes. The y-axis represents the log-transformed estimated effect sizes from the simulated data. The spearman correlation coefficient between these values using the BIRDbath model is 0.503, and the mean squared error of the estimate is 0.057. The spearman correlation coefficient between these values using the BIRD model is 0.357, and the mean squared error of the estimate is 0.297. (b-c) Representations of decreasing error of estimation and increasing correlation when dividing data among different pools across the allele frequency spectrum. The data are separated into pools (red) and collapsed into a single pool (blue). The X-axis represents the maximum minor allele frequency of the variants. The Y-axis represents (d) the MSE of the estimated effect when compared to the simulated effect, and (e) the spearman correlation between the estimated and simulated effects. (d-e) Representations of decreasing error of estimation and increasing correlation when dividing the data into additional pools. The X-axis represents the number of pools in the simulation. The Y-axis represents (d) the MSE of the estimated effect when compared to the simulated effect, and (e) the spearman correlation between the estimated and simulated effects.

### 2.3 Structured STARR-seq on 100 Thousand Genomes Project individuals

The 100 genomes assayed in the pooled STARR-seq experiment together contained ∼27 million genetic variants across the genome, including 99.5% of common (as defined by gnomAD allele frequencies^24^) variants in the full cohort of Thousand Genomes samples spanning 2,504 individuals, as well as 90.1% of uncommon, 41.2% of rare, and 5.3% of ultra-rare variants in the entire Thousand Genomes cohort. 4.2% of the variants in this study had unknown allele frequencies in the general population (Fig. 4a). The fragments in these libraries were on average 435 bp (Fig. 4b), comparable in size to a sample of caQTL open chromatin regions from the AFGR^22^. Each individual pool includes on average ∼1 billion unique DNA fragments, covering the genome at ∼150X on average (Fig. 4c). Mean coverage within STARR-seq and ATAC-seq peaks was 24, where some peaks and some pools had better coverage than others (e.g., Fig. 4d; Table 1).

**Figure 4.**
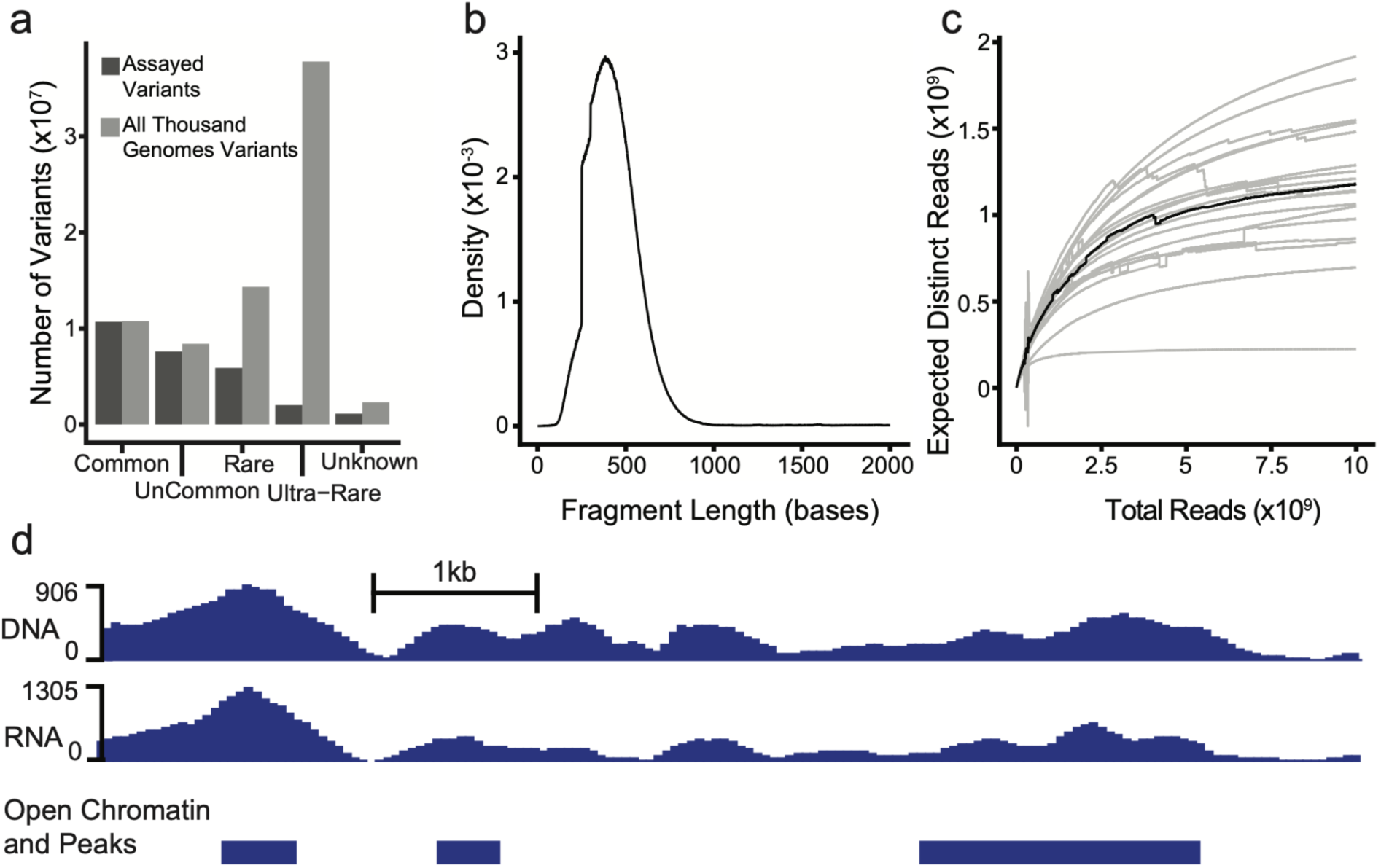
(a) Density plot of the fragment length distribution of the DNA fragments. X-axis represents the fragment length. Y-axis represents the proportion of reads that exist at that fragment length. Individual replicates of pools are represented in gray, and an averaged curve is represented in black. (b) Library complexity plot for the input libraries. X-axis represents the total reads sequenced. Y-axis represents the number of distinct reads at the corresponding sequencing depth. Individual estimates for each replicate of each pool are represented in gray, and an averaged curve is represented in black. (c) Allele frequency distributions for different categories of allele rarities. X-axis represents the different allele rarity categories. Y-axis represents the count of variants in each category. Common alleles have minor allele frequencies greater than 0.05, uncommon alleles have minor allele frequencies between 0.01 and 0.05, rare alleles have minor allele frequencies between 0.01 and 0.001, and ultra-rare variants have minor allele frequencies less than 0.001. Allele frequencies are taken from the GnomAD database. The light gray bars represent all variants present in the Thousand genomes project. The dark gay bars represent the variants present in the 100 individuals included in this study. (d) A browser track of the ATG10 locus in chromosome 5. X-axis represents genomic coordinates. Y-axis represents the number of reads overlapping each position.

Table 1.

Table 1
| Pool | Average DNA coverage | Average RNA coverage |
| --- | --- | --- |
| 1 | 125 | 86 |
| 2 | 76 | 144 |
| 3 | 78 | 138 |
| 4 | 5 | 14 |
| 5 | 11 | 2 |
| 6 | 12 | 26 |
| 7 | 12 | 41 |
| 8 | 12 | 31 |
| 9 | 10 | 21 |
| 10 | 12 | 18 |
| 11 | 16 | 25 |
| 12 | 13 | 26 |
| 13 | 11 | 27 |
| 14 | 14 | 29 |
| 15 | 13 | 20 |
| 16 | 13 | 27 |
| 17 | 11 | 23 |
| 18 | 11 | 19 |
| 19 | 12 | 33 |
| 20 | 13 | 26 |

We used the BIRDbath model to estimate regulatory effects of variants in these data. In total, we assayed the effects of 1,520,879 biallelic variants in peak regions across all 20 pools. Most variants had an estimated effect close to zero, with a median of 0.01863 (Figure 5a), and the majority of variants had low posterior probability for the variant to have an effect more extreme than our null threshold (log fold-change of 1.25, in either direction) (Figure 5b).

**Figure 5.**
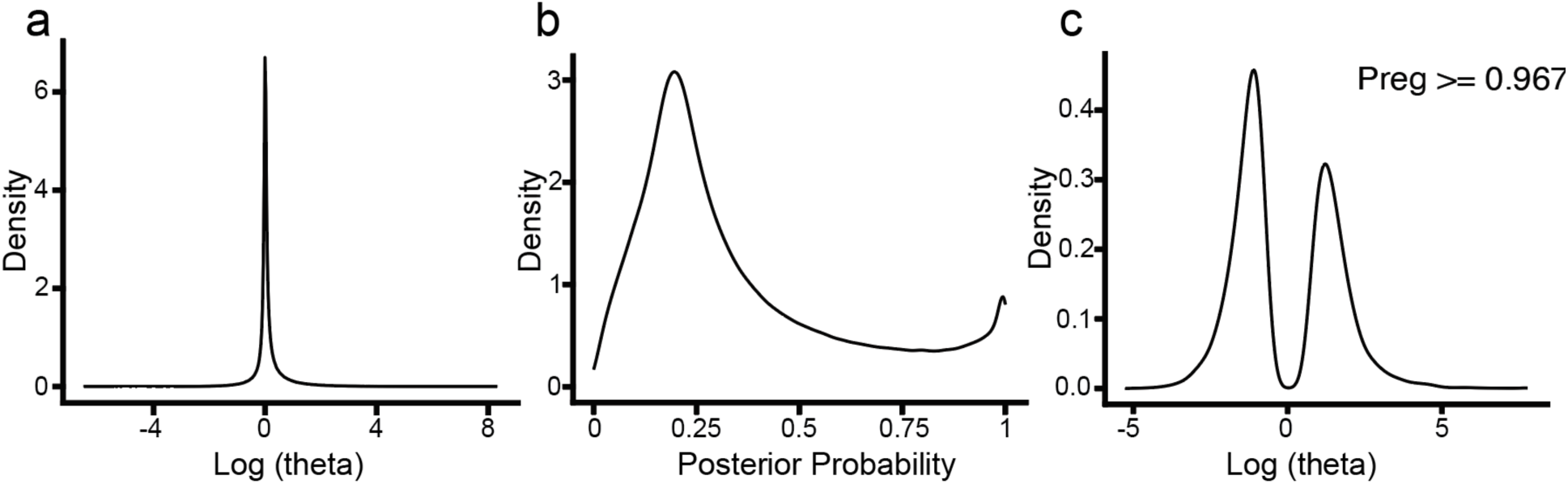
(a) Distribution of effect sizes of all variants as estimated by BIRDbath. X-axis represents the log-transformed effect size. Y-axis represents the proportion of variants with that effect size. (b) Density plot of the distribution of posterior probability that a STARR-seq variant will have an effect. X-axis represents the posterior probability (P_reg_). Y-axis represents the proportion of variants at that probability of having an effect. (c) Density plot of the distribution of the effect sizes of significant variants as they were estimated by the BIRDbath model. Here, significance is defined as a variant having a posterior probability of having an effect above the 95^th^ percentile (posterior >= 0.967). X-axis represents the log-transformed effect size. Y-axis represents the proportion of variants with that effect size.

Together, these results indicate that most variants do not have a regulatory effect, consistent with our expectations. Of the significant variants (posterior probability greater than the 95^th^ percentile), estimated effects had a mean magnitude of ∼1.4883 +/− 0.6839 SD and they were slightly more likely to have a negative effect than a positive one (Figure 5c).

We compared estimated STARR-seq variant effects to estimated changes to transcription factor (TF) affinity for known DNA TF motifs using motifbreakR. At an FDR-corrected significance of 0.05, the concordance between STARR-seq effect and TF motif effect was significant for 58 TF clusters^25^. The three most significant concordances were for AP1, ETS, and CREB (Figure 6a), all of which are known to be expressed and known to be transcriptional activators in K562. The more conserved positions in the PWM are more likely to have associated STARR-seq variants, as indicated by their information content (Figures 6b, c).

**Figure 6.**
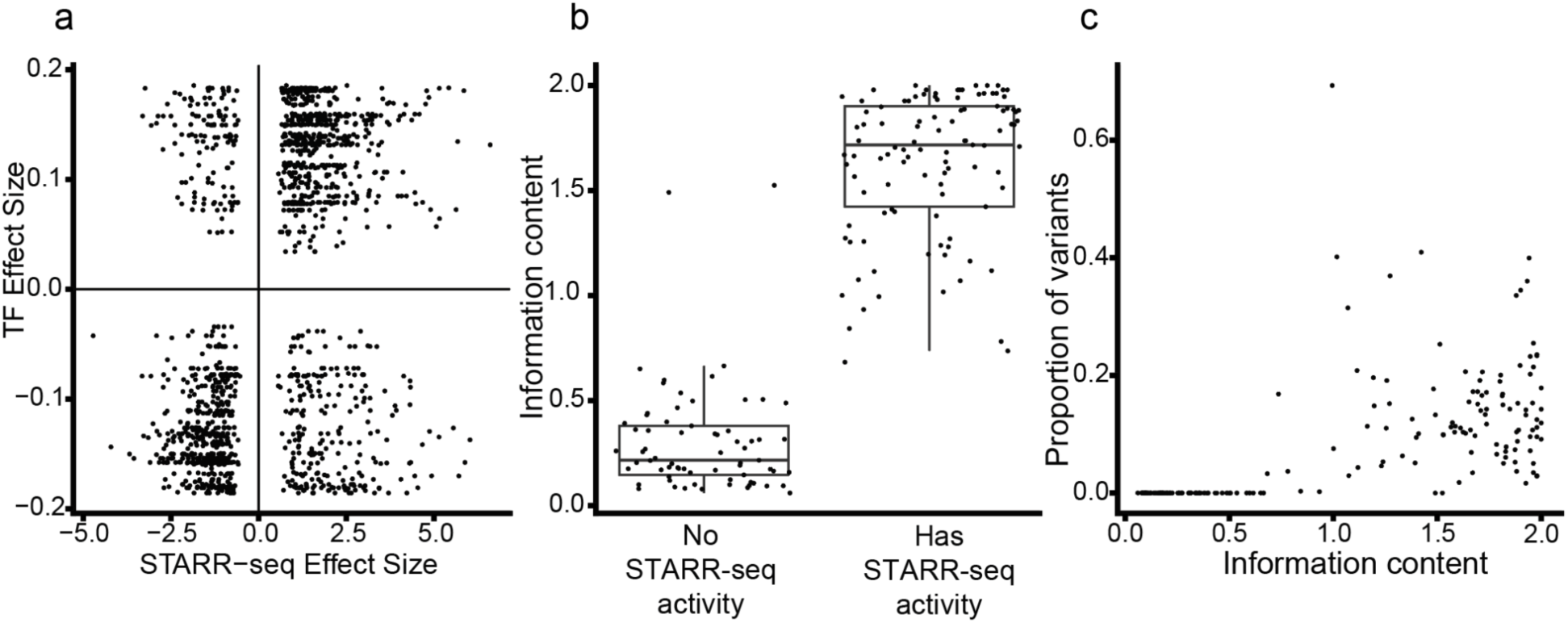
(a) Concordance of magnitude and direction of STARR-seq effects and estimated effects caused by changes to transcription factor binding motifs (AP-1, CREB, ETS). Each point represents a different variant in a concordant transcription factor motif. X-axis represents the effect size of the variant in the STARR-seq assay. Y-axis represents the estimated change in transcription factor binding. With these three TFs, this concordance is significant (Fisher’s exact test: p=3.094601e-84; Chi-squared test: p=7.644794e-80) (b) Difference in information content of transcription factor positions with and without STARR-seq activity. Each point represents a different position in the motifs. X-axis represents the categories of having or lacking STARR-seq activity. Y-axis represents the information content at each position. These categories are significantly different (t-test: p=3.103872e-67). (c) Trend in the comparison between the proportion of variants at each position in a motif relative to the information content at that position. Each point represents a different position in the motifs. X-axis represents information content at that position. Y-axis represents the proportion of variants that exist in that position compared to all variants in that motif.

We compared results of the STARR-seq experiments to the results of prior GWAS experiments. Here, we compare to the African Functional Genomics Resource (AFGR)^22^, which provides expression and chromatin accessibility QTLs found in a population of African individuals. This dataset was selected because all samples included in our STARR-seq assays were from African ancestries as well, so the ancestry between the two datasets match.

First, we compared the directional effect sizes of the STARR-seq variants and the QTLs that exist within the same open chromatin region. For each QTL, we aggregated the effects of all significant STARR-seq variants in an associated open chromatin region, accounting for phasing and LD. We found 70% concordance between the STARR-seq data and the caQTLs (Figure 7a), and 66% concordance between the STARR-seq data and the eQTLs (Figure 7b).

**Figure 7.**
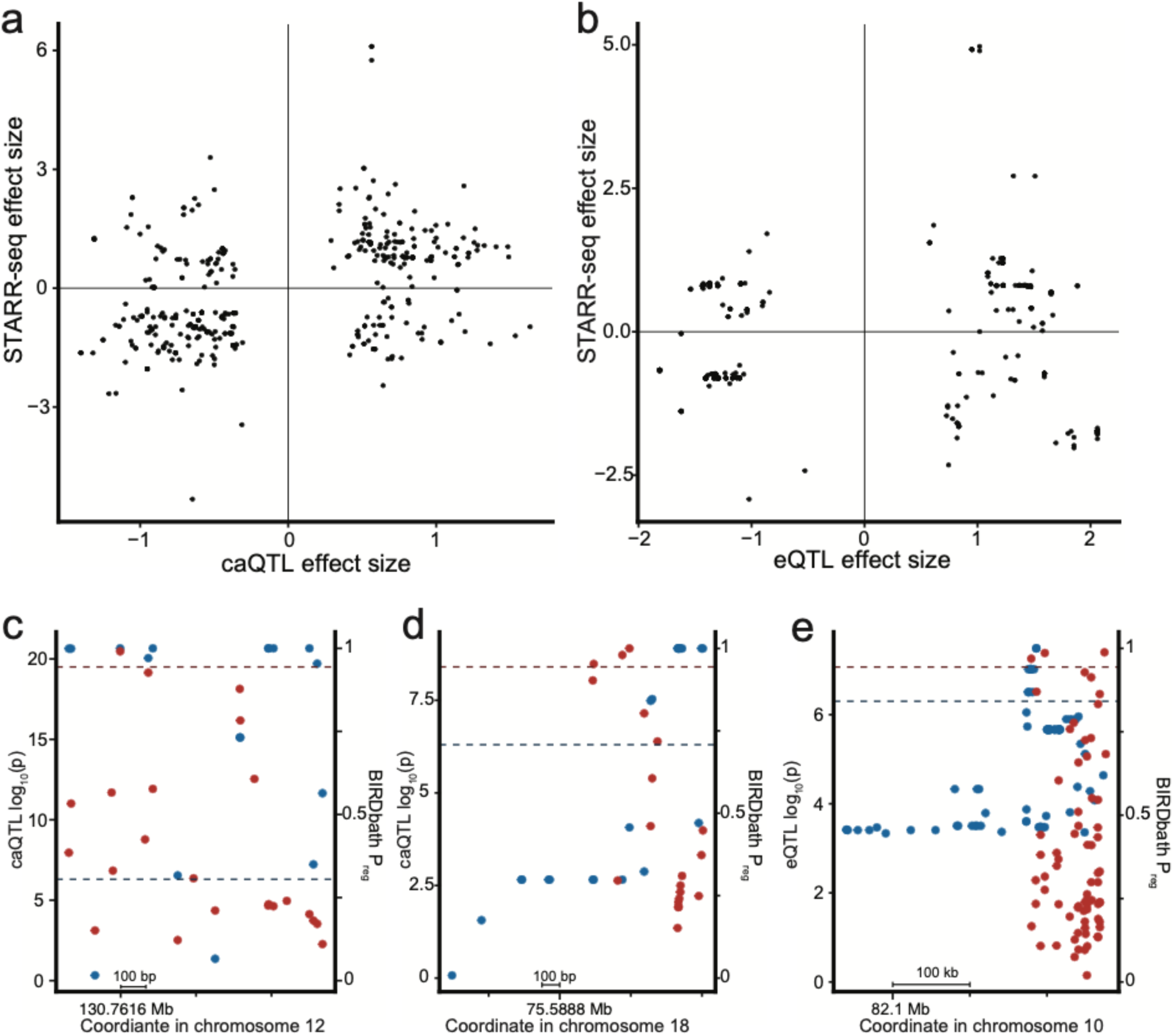
(a-b) Concordance plots showing the magnitude and directions of STARR-seq effects and (a) caQTL or (b) eQTL effects. Each point represents a different cis-QTL with at least one STARR-seq variant in associated open chromatin regions. The X-axis represents the effect size of the QTL. The Y-axis represents the aggregated effect sizes of all STARR-seq variants in the associated open chromatin region. When comparing to caQTLs, these datasets have a concordance of 70% (fisher’s exact p = 9.775046e-17, chi-squared p = 2.959211e-16), and a Pearson correlation of 0.385 (p=5.216174e-16). When comparing to eQTLs, these datasets have a concordance of 66% (fisher’s exact p = 9.326762e-10, chi-squared p = 1.903943e-9), and a Pearson correlation of 0.187 (p=1.94745e-4). (c-e) Functional annotation figures comparing the resolution of significant QTLs and STARR-seq variants. The X-axis represents the chromosomal coordinate. Blue points are QTL p-values, represented on the left Y-axis. Red points are STARR-seq P_reg_ values, represented on the right Y-axis. The regions depicted with caQTLs are (c) chromosome 12 from position 130,761,358 to 130,762,412 and (d) chromosome 17 from position 75,588,068 to position 75,589,662. The region depicted with eQTLs is (e) chromosome 5 from position 82,279,462 to position 82,386,977.

Using these open chromatin regions, we assessed the ability of the STARR-seq variants to functionally annotate QTLs in these regions. Here, we compared the –log10 p-value of the QTLs to the Preg values of the STARR-seq variants which came from BIRD. We repeated this analysis with both the caQTL dataset and the eQTL dataset.

We found 216 open chromatin regions with caQTLs which overlap with STARR-seq variants. Of those, 52 had enough variants present to draw conclusions, and they fell into four main categories. 17 had multiple lead STARR-seq variants (Figure 7c, Suppl. Fig. 6), 12 had a lead STARR-seq variant which overlapped a lead QTL (Fig. 7d, Suppl. Fig. 7), 11 had a lead STARR-seq variant which overlapped a non-significant QTL (Suppl. Fig. 8), and 12 had a lead STARR-seq variant which was missing from the QTL dataset (Suppl. Fig. 9). We repeated this with the eQTL dataset. All 29 open chromatin regions had sufficient QTLs and STARR-seq variants. There were 14 regions where there were multiple significant STARR-seq variants in the open chromatin region (Fig. 7e, Suppl. Fig. 10), 2 regions where the lead STARR-seq variant overlapped a significant QTL (Suppl. Fig. 11), and 13 regions where the lead STARR-seq variant did not overlap a QTL (Suppl. Fig. 12).

## 3. Discussion

In this study we proposed and evaluated a novel experimental design for population-scale high-throughput reporter assays, and characterized the properties of that design with respect to its ability to reveal measurable noncoding variant effects.

Using a cohort of 100 individual samples from the Thousand Genomes Project, we demonstrated that segregating samples into distinct pools results in a substantial increase in accuracy of estimates of variant effect, likely due to the inflation of allele frequencies within individual pools as compared to a non-structured or “collapsed” design in which a single plasmid library is constructed for the entire cohort. That difference was largest for rare variants, which are often the variants of greatest interest in disease studies and often under the strongest negative selection, and which are also the variants for which estimation of effect size tends to be most under-powered^23^.

Indeed, a prime motivation for the partitioning of samples into distinct pools is to mitigate dropout of individual alleles in pooled multi-sample experiments. Dropout--when an element or variant present in the original sample material is not present in the resulting experimental readout--can occur at various points in the experiment, from capture and integration of genomic DNA fragments into plasmids, to PCR amplification, to high-throughput sequencing and post-processing of raw data. As a crude illustration of the potential for this approach to reduce dropout during the plasmid library construction, note that under simple binomial sampling of fragments at 150X coverage, the probability of an individual dropout event for a non-structured cohort of 100 diploid individuals is P(x=0 | N=150, p=1/200) = 0.47, or nearly 50% allelic loss per site, while for our structured design with 20 pools that probability is reduced by several orders of magnitude to P(x=0 | N=150, p=1/10) = 1.4e-7.

Note, however, that dropout during sequencing remains a risk for low sequencing coverage. In this study our sequencing depth was ∼7 reads on average within peaks, so that our expected dropout rate during sequencing, given a mean allele frequency of 0.1, was P(x=0 | N=7, p=0.1) = 0.48 under simple binomial sampling, and indeed, we observed a total rate of ∼22% dropout of heterozygous sites in the plasmid library as compared to expected from genotyping (20913226 heterozygous sites were detected in the DNA sequenced from the plasmid library, out of 26792719 genotyped as heterozygous in our cohort), which may include both dropout during library construction and sequencing. As our proposed experimental design does not address dropout due to sequencing depth, it is important to ensure adequate sequencing coverage to reap the full benefits of this assay.

The structured pooling design also produces a richer readout that facilitates more precise estimation of effect sizes and greater power to detect and annotate functional regulatory variants. That improvement can be explained by changes in allele frequencies brought about through the partitioning of samples. As noted above, the largest improvements in estimation accuracy were seen for variants with the lowest allele frequencies in the original cohort. For rare variants that are present in only one pool, the allele frequency in that pool will be substantially larger in our design than in the whole cohort, possibly explaining the larger improvement in estimation accuracy among rare variants. For less rare variants that occur in multiple pools, partitioning will generally produce greater heterogeneity in allele frequencies as compared to a collapsed structure having only one, global pool.

As we have shown previously, that heterogeneity in allele frequencies results in a Poisson-binomial sampling process that reduces variance in allelic read counts, and empirically was found to reduce the mean error in effect estimates. That resulting improvement in effect estimates is seen even with a simple estimator based on ratios of read counts, without requiring specialized modeling. In the present work we have shown that additional gains in estimation accuracy can be accrued by explicitly modeling that heterogeneity. In direct comparisons between the original BIRD model and our updated model, BIRDbath, the new model was seen to produce significantly more accurate effect estimates through the explicit modeling of per-pool allele frequencies and the heterogeneity of those frequencies between pools.

The value of high-throughput reporter assays for functional annotation of regulatory variants has been well demonstrated previously^20,23^. In the present study we added to the body of evidence that these assays can produce outputs that are concordant with other indicators of likely noncoding functionality, including both caQTLs and eQTLs, as well as signals of selection on individual nucleotide positions within TF binding motifs. These results support the importance of continued improvement to both assay design and methods for statistical analysis of data produced by those assays.

The structured pooling design described here incurs greater costs in both labor and experimental materials, even when sequencing depths are kept fixed, by requiring the construction of multiple plasmid libraries. With the specific design employed in this study, we have attempted to strike a balance in the tradeoff between cost and utility. While the single-library “collapsed” design minimizes effort for library construction, the other extreme of constructing a plasmid library for each individual in a large cohort would produce much richer data by elevating within-pool allele frequencies, but would be prohibitively expensive in the limit. There thus remains a need for bioinformatic tools to help explore the range of options in granularity of the pooling structure so as to optimize the cost-benefit tradeoff in terms of improved variant annotation versus experimental cost. Variant effect models such as the one provided in this study can be readily used in the construction of power analysis tools to aid in those experimental design decisions.

A related future direction for this work lies in developing methods for optimally assigning individual samples to experimental pools. As this assignment directly impacts allele frequencies within pools, the accuracy of variant effect estimates for an individual variant may be substantially impacted by that assignment. However, because sample assignments impact all variants jointly, and because the optimal assignment for one variant may be different from the optimal assignment for another variant, the problem of optimal assignment across all variants jointly may be nontrivial, and may depend in complex ways on patterns of shared ancestry and possibly haplotype sharing between individuals in the cohort. While our choice of African samples in this study was aimed at maximizing overall genetic diversity, the question of which individuals from a population to include in an experimental cohort to maximize informativeness of these assays also remains a relevant and unanswered question for future research.

## 4. Materials and Methods

### 4.1 A realistic simulator of structured STARR-seq experiments

To test the accuracy of the BIRDbath model, a simulator was created to generate synthetic STARR-seq data (Fig. 8), using a graphical model to represent conditional distributions of variables to simulate based on empirical measurements. To maximize the realism of the simulations, they were based on real data from the first pool in our 20-pool experiment. In particular, genotype frequencies and effect sizes were sampled from their joint empirical distribution, with empirical effect sizes being defined by their estimates from our previous model applied to read data from pool 1^23^. In this way, the empirical relationship between the population allele frequency and the effect size of the variant is maintained. Conditional on those effect sizes and total counts, binomial sampling was then used to draw individual allelic read counts that served as inputs for testing the BIRDbath model. Additional details are provided in Suppl. Text. S1.

**Figure 8.**
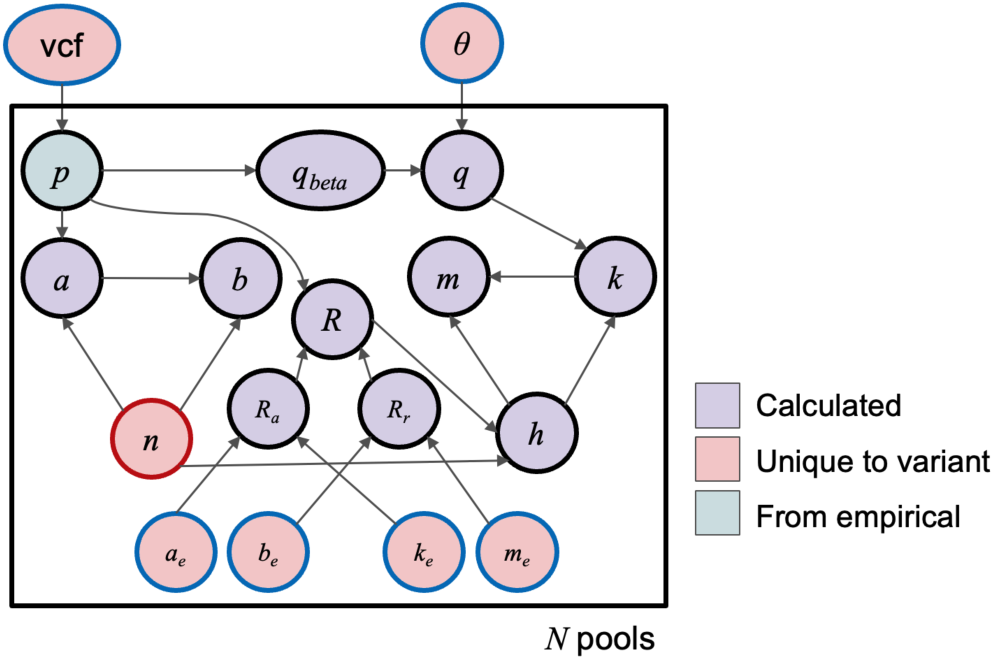
Graphical representation of a simulator for STARR-seq data. See text for variable definitions. Each variable was sampled conditional on each of its parent variables in the graph.

The result of this simulation is a dataset of synthetic STARR-seq data with a variety of allele frequencies and effect sizes for the simulated variants, along with a list of effect sizes that were used to generate allele count data for each of the simulated variants. The simulated dataset included experimental data for 20 pools of 5 individuals, with one replicate of each pool.

### 4.2 Assessing the impact of pool heterogeneity on effect estimates

The simulator described in section 4.1 was modified to test the impact of heterogeneity in allele frequency among pools and replicates. Unlike the simulator described in section 4.1, which simulated exactly 5 genomes per pool, arbitrary numbers of genomes and pools are simulated here in order to address heterogeneity arising stochastically during pool or replicate construction. Specifically, 100 genotypes were randomly selected from all of the Thousand Genomes Project samples to be included in these simulations. These genotypes were divided into any number of experiments between 1 and 100.

Simulated allele counts were then sampled from a binomial distribution based on those genotypes. Additionally, the total coverage of the simulated data was held constant at 1000 to control for differences in sequencing depth while varying other parameters. The coverage in a single pool was calculated based on the simulations described in Majoros et.al. 2020.

A dataset which included replicates of a single pool was simulated for comparison. Here, the samples were divided in the same way as when simulating pools, and one of the pools was randomly selected to be repeated as a replicate (but with counts simulated anew from the same frequencies). One replicate of DNA and multiple replicates of RNA were simulated. Total coverage was again held constant across all replicates of simulated RNA libraries.

For these analyses, in which no model is used, the effect size values are calculated using a simple method we refer to as the *naive estimator*:

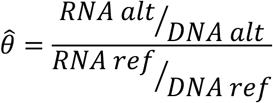

The RMSE of the estimate was defined as the RMSE between the estimated and simulated true theta.

### 4.3 Graphical model and inference

BIRDbath is a modified version of BIRD^23^, with the primary differences being that read count variables are grouped by pool in addition to replicate (Figure 9), and the known DNA alternate allele frequencies from DNA genotyping are provided to the model in variable ω_2_, which serves as the mode of a beta prior *p_iz_* ∼ *beta*(ω*_z_*, *c*_2_) parameterized via mode and concentration^26^. Experiments with only one replicate per pool are naturally supported.

**Figure 9.**
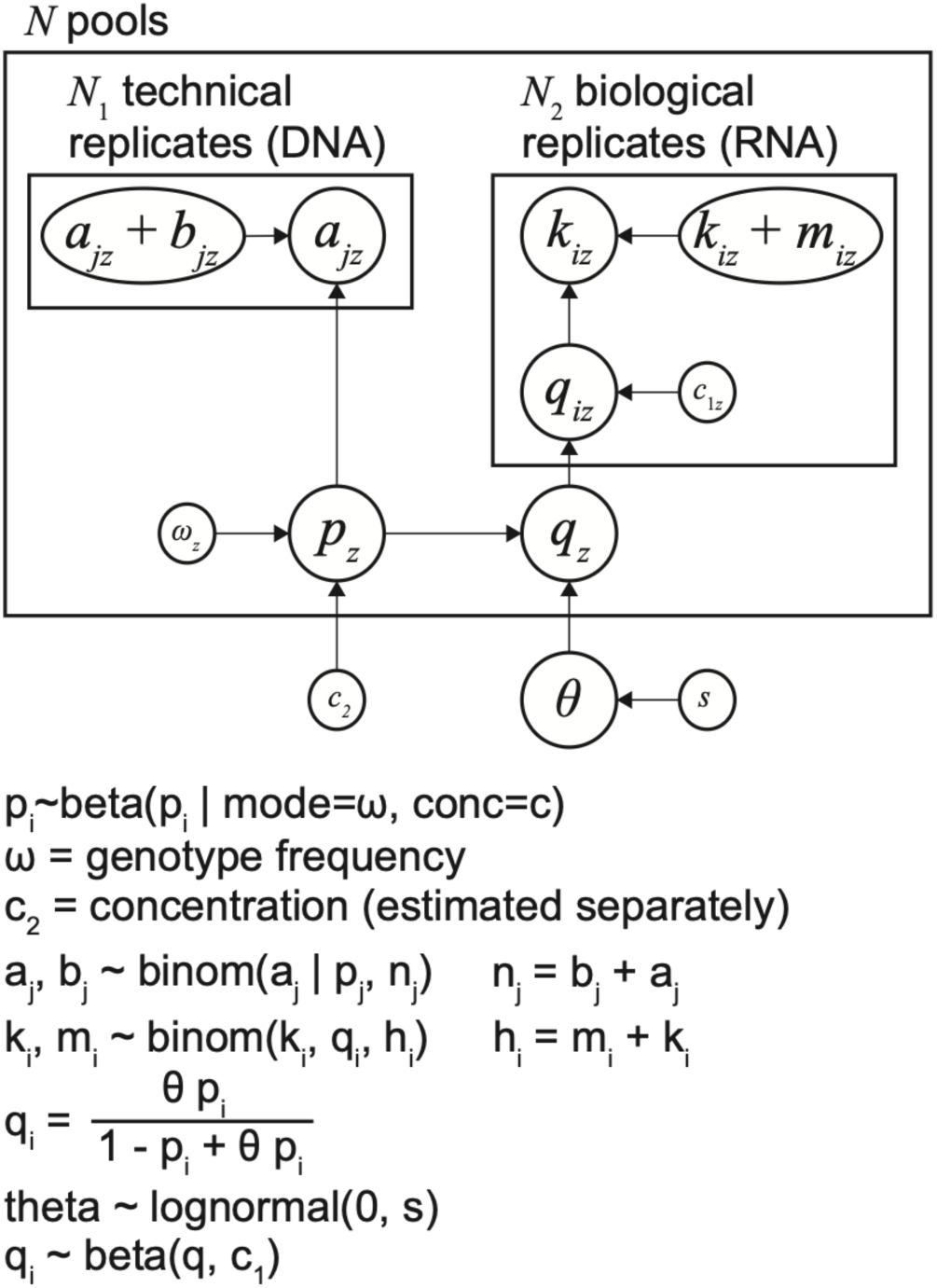
Probabilistic graphical model for BIRDbath. Nodes represent observed or latent variables, and arrows denote conditional dependence relations. Boxes represent replication of all variables within each box across replicates or pools, as indicated by variable subscripts. See Table 1 for variable definitions.

As in BIRD, STAN is used for inference^27^. In short, the graphical model (Fig. 9, Table 2) denotes a dependency structure and the joint distribution of all variables in the model is factored according to that structure, as a product of likelihoods and priors. The latter terms are used within STAN to construct an acceptance ratio as employed in the Metropolis-Hastings algorithm^28,29^. STAN proposes samples via a Hamiltonian Monte Carlo proposal distribution and accepts or rejects according to the aforementioned ratio, producing a chain of MCMC (Markov chain Monte Carlo) samples. 1000 warmup samples are drawn, which are discarded, followed by 1000 additional samples which are retained and used to produce a posterior median estimate of the effect size, θ. BIRDbath reports the posterior median estimate as well as a 95% credible interval estimated from the MCMC chain. In addition, for a given ignorance interval (also known as a region of practical equivalence or RoPE^26^), BIRDbath reports the posterior mass outside of that interval, referred to as *P_reg_*.

**Table 2:** Definitions of variables in Fig. 8.

| Variables | Definitions |
| --- | --- |
| $p_z$ | Alternate allele frequency in plasmids of the $z^{\text{th}}$ pool (DNA) |
| $q_z$ | Alternate allele frequency in RNA of the $z^{\text{th}}$ pool |
| $q_{iz}$ | Alternate allele frequency in $i^{\text{th}}$ RNA replicate of the $z^{\text{th}}$ pool |
| $\theta$ | Effect size of the variant |
| $a_{jz}$ | Alternate allele count in DNA sequencing reads from $j^{\text{th}}$ replicate of the $z^{\text{th}}$ pool |
| $b_{jz}$ | Reference allele count in DNA sequencing reads from $j^{\text{th}}$ replicate of the $z^{\text{th}}$ pool |
| $k_{iz}$ | Alternate allele count in RNA sequencing reads from $i^{\text{th}}$ replicate of the $z^{\text{th}}$ pool |
| $m_{iz}$ | Reference allele count in RNA sequencing reads from $i^{\text{th}}$ replicate of the $z^{\text{th}}$ pool |
| $c_{1z}$ | Concentration parameter for $q_{iz}$ |
| $c_2$ | Concentration parameter for $p_z$ |
| $\omega_2$ | Genotype frequency, expected value for $p_z$ |
| $s$ | Variance parameter for prior on theta |

BIRDbath accounts for the nested structure of replicates within pools in the new experimental design for structured STARR-seq. The nesting structure of the model allows for different numbers of replicates per pool, as well as different replicate structures for RNA compared to DNA. Importantly, DNA and RNA replicates are not assumed to be paired, as in practice DNA replicates are typically technical replicates and RNA are typically biological replicates.

Sampling statements:

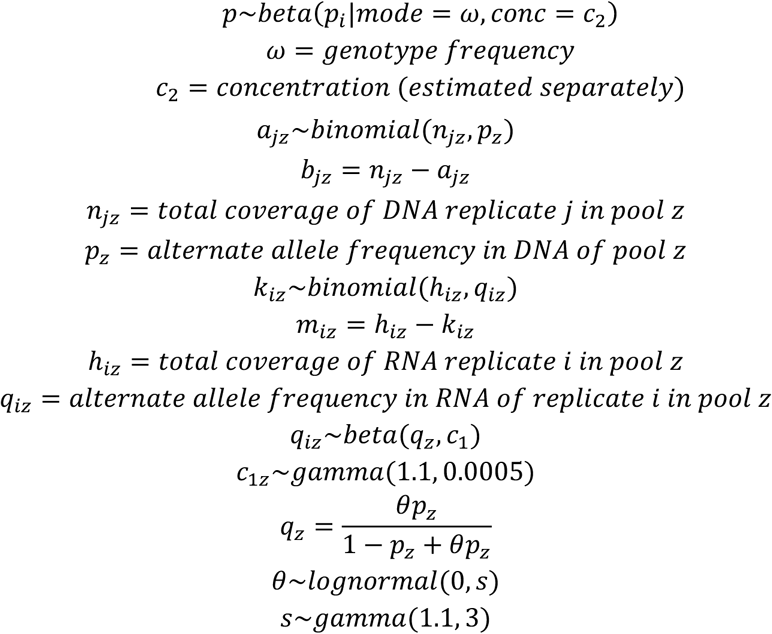

### 4.4 Model benchmarking

To test the accuracy of the BIRDbath model, the simulator described in 4.1 was used to simulate 20 pools of 5 individuals for input to the model. Inference was performed (section 4.5) to produce a chain of MCMC samples from the posterior distribution for theta. The posterior median of that chain was used as the point estimate of the model for comparison to the true simulated value. MSE and Spearman correlation (rho) were reported for these comparisons.

Assessment of model accuracy across the minor allele frequency spectrum was based on the minor allele frequencies of the variants in the overall set of simulated variants.

Allele counts were collapsed by summing the simulated counts of alleles across pools in each category separately: DNA reference, DNA alternate, RNA reference, and RNA alternate.

First, the BIRDbath model was used on the simulated data with 20 pools of 5 individuals to determine effect sizes from the entire dataset. This dataset is considered “structured” because it includes the information about each pool individually. Then, the allele counts in each library of each pool were summed to get overall allele counts as if the entire series of simulated experiments were done as a single replicate of one pool. This method of summing all of the allele counts is referred to as “collapsing”, and it results in the different organization of the structured and collapsed versions of the datasets.

Because the data were simulated with known effect sizes, the effect sizes that have just been estimated using the BIRDbath model can be compared to those that were used in the simulation of the data. The mean squared error between these two values and the Spearman correlation were used to assess the relative accuracy of these models.

To assess the impact of pooling structure on the accuracy of variant effect estimation, the structured dataset with 20 pools of 5 individuals was re-structured. The number of pools was changed to 10 pools of 10, 5 pools of 20, 4 pools of 25, 2 pools of 50, and finally 1 pool of 100. When decreasing the number of pools, the original 20 pools were grouped into equal sized sets. The counts of alleles were collapsed within this set for each variant to create a new aggregated pool with all variants, for each of the sets individually. BIRDbath was used to estimate effects for all of the variants in each restructured dataset individually. As above, the mean squared error and Spearman correlation of the effect size used in the simulation and the effect size estimated with BIRDbath were used to assess the relative accuracy of these models.

No new datasets were generated to assess the impact of pooling on different rarities of variants. The original structured dataset with 20 pools of 5 individuals was used. The dataset was divided into 10 groups depending on the minor allele frequency in the study population, where the maximum minor allele frequencies were multiples of 0.05. These subsets were:

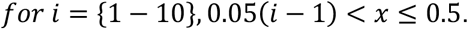

Once again, each subset of the dataset was assessed with the mean squared error and the Spearman correlation of the effect size used in the simulation and the effect size estimated with BIRDbath.

### 4.5 Application to genomic data

In order to apply BIRDbath to genomic data, all sequencing samples from the STARR-seq assay must be aligned, and the occurrences of each variant in each replicate of the DNA and RNA libraries must be counted using a tool such as samtools mpileup^30^. The input file for BIRDbath is generated from the data from these count files, the expected alternate allele frequencies in each pool for each variant, and the pool size. To simplify the generation of BIRDbath files, wrapper scripts are provided at https://github.com/bmajoros/BIRDbath.

### 4.6 Experimental methods

The STARR-seq assay was performed following the methods in Dura <u>et.al</u> 2025. We designed these assays using a structured pool STARR-seq approach, separating the samples into 20 pools of 5 and performing STARR-seq on each pool, and sequencing different numbers of replicates for each library of each pool (Table 3).

**Table 3:** List of Thousand Genomes samples included in each pool.

| Pool Number | Sample List | DNA replicates | RNA replicates |
| --- | --- | --- | --- |
| 1 | NA18517, NA18519, NA18858, NA18868, NA18873 | 3 | 3 |
| 2 | NA18910, NA19149, NA19236, NA19247, NA19256 | 6 | 16 |
| 3 | NA18488, NA18499, NA18508, NA18510, NA18870 | 3 | 8 |
| 4 | NA18502, NA18867, NA18907, NA18933 NA19102 | 1 | 5 |
| 5 | NA19099, NA19107, NA19114, NA19119, NA19121 | 2 | 8 |
| 6 | NA19093, NA19118, NA19137,<br>NA19138, NA19210 | 1 | 1 |
| 7 | NA19098, NA19113, NA19152,<br>NA19171, NA19190 | 1 | 1 |
| 8 | NA19144, NA19207, NA19214,<br>NA19222, NA19257 | 1 | 1 |
| 9 | NA18505, NA18909, NA18916,<br>NA18923, NA19175 | 1 | 1 |
| 10 | NA19096, NA19116, NA19130,<br>NA19159, NA19197 | 1 | 1 |
| 11 | NA18511, NA19147, NA19185,<br>NA19200, NA19248 | 1 | 1 |
| 12 | NA18489, NA18908, NA18934,<br>NA19095, NA19184 | 1 | 1 |
| 13 | NA19153, NA19160, NA19172,<br>NA19189, NA19223 | 1 | 1 |
| 14 | NA19108, NA19141, NA19146,<br>NA19204, NA19235 | 1 | 1 |
| 15 | NA18486, NA18498, NA18520,<br>NA18912, NA19209 | 1 | 1 |
| 16 | NA18861, NA18917, NA19198,<br>NA19201, NA19206 | 1 | 1 |
| 17 | NA19092, NA19117, NA19129,<br>NA19131, NA19143 | 1 | 1 |
| 18 | HG02107, HG02462, HG02464,<br>NA19213, NA19225 | 1 | 1 |
| 19 | HG02594, HG02611, HG02623,<br>HG02635, HG02645 | 1 | 1 |
| 20 | HG02666, HG02771, HG02804,<br>HG02811, HG02881 | 1 | 1 |

### 4.7 Statistical analyses of experimental data

To assess population allele frequencies of the assayed variants, population frequency values were queried from the gnomAD database.

To ensure that variants included in the analysis had regulatory activity, variant effects were only assessed within these peak regions. Peaks were called using MACS to determine individual peak regions in each pool, as well as peaks found when aggregating all data into a single file. This was combined with a region of ATAC-seq peaks^31^.

Changes to transcription factor motif binding were estimated using the R package motifbreakR for all PWMs in the HOCOMOCO database. Motifs assigned to a single variant were down-sampled to include only one motif within a pre-defined motif cluster^25^. When down-sampling, the specific PWM that was chosen was the one with the greatest match score to the sequence surrounding the variant, regardless of if that score was assigned to the reference or alternate allele. The effects estimated from BIRDbath for each variant were compared to those that were found with motifbreakR. A motif cluster was defined as being concordant if the direction of the transcription factor binding effect from motifbreakR and the allelic effect from BIRDbath were in the same direction at least 70% of the time, and the motif had to have at least 500 associated variants.

QTLs were correlated with aggregated haplotype effect sizes from the BIRDbath model. To aggregate BIRDbath effect sizes of individual variants across a haplotype, first a phased LD score had to be calculated between the QTL and all STARR-seq variants located in an associated open chromatin region. The aggregated effect size was represented by the sum of the products of the STARR-seq effect size, and the phased LD score for each variant in the open chromatin region.

## Data and Software Availability

Accessions for raw data in ENCODE portal: https://www.encodeproject.org/functional-characterization-experiments/ENCSR926NDZ/

Accessions for raw data & estimates in IGVF portal:

/analysis-sets/IGVFDS4714VWGB/

/analysis-sets/IGVFDS6219XNKZ/

/analysis-sets/IGVFDS6954KEHC/

/analysis-sets/IGVFDS5833SGWM/

/analysis-sets/IGVFDS5241INPJ/

/analysis-sets/IGVFDS8554TSTG/

/analysis-sets/IGVFDS9219WLAH/

/analysis-sets/IGVFDS2766BVRH/

/analysis-sets/IGVFDS2175LLDQ/

/analysis-sets/IGVFDS9967RCHV/

/analysis-sets/IGVFDS5400TMWO/

/analysis-sets/IGVFDS1147GIJW/

/analysis-sets/IGVFDS3299RYZX/

/analysis-sets/IGVFDS6245HBJO/

/analysis-sets/IGVFDS6846CJYR/

Source code repository: https://github.com/bmajoros/BIRDbath

## Author contributions

Experimental data generation: Keith S, Kari S, SM, CZ

Model and software development and analysis: KD, WHM

Conceptualization and supervision of experimental work: TER

Conceptualization and supervision of computational work: WHM, TER, ASA

## Acknowledgements and Funding

Research reported in this publication was supported by the National Institute of General Medical Sciences (NIGMS) of the National Institutes of Health (NIH) under award number 1R35-GM150404 to W.H.M., and National Human Genome Research Institute (NHGRI) of NIH 5U01-HG011967 to A.S.A. and W.H.M., and NHGRI 5UM1-HG012053 to T.E.R. Content is solely the responsibility of the authors.

## Notes

### Competing Interest Statement

The authors have declared no competing interest.

