## Supplementary text and figures for "Structured Pooling Improves Detection of Rare Regulatory Mutations in Population-Scale Reporter Assays"

### Supplementary Figures

#### Supplementary Figure 1

a

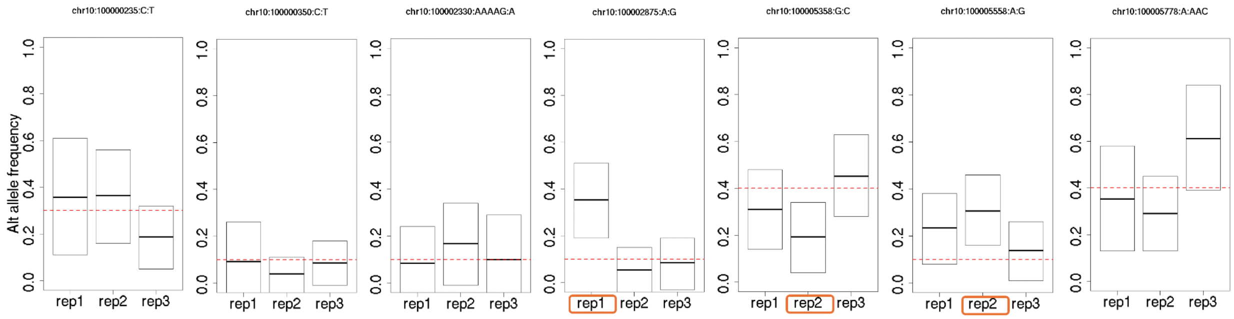


b

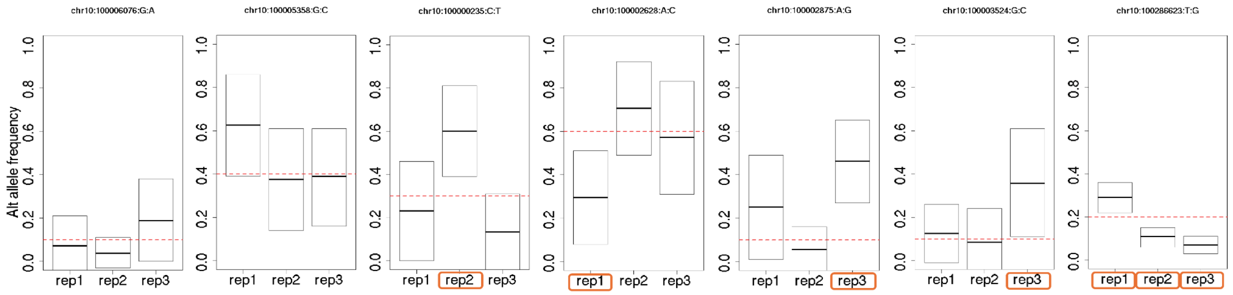


Heterogeneity in the experimental data was assessed by comparing the allele frequencies of individual variants to the expected range of allele frequencies given binomial variance.

Supplementary Figure 2

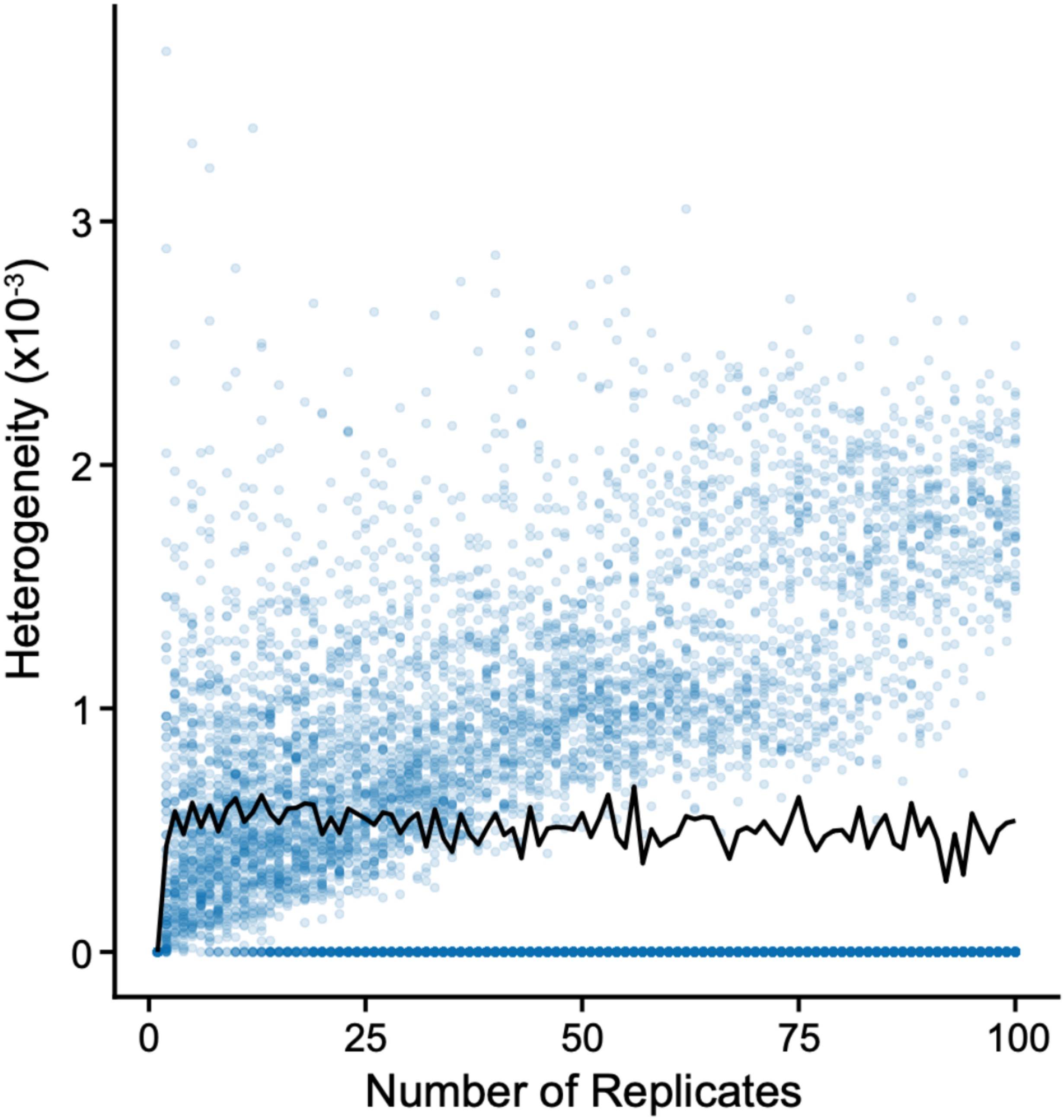


Heterogeneity is represented as a function of the number of replicates of a single experiment. In this case, there are 100 total samples assayed across all replicates. The number of replicates is represented on the X-axis. Heterogeneity in RNA is defined as the sample variance in the alternate allele frequency in the RNA, represented on the Y-axis. The black trend line indicates the average heterogeneity at each number of replicates assayed. Homozygous cases, where all samples in the pool being replicated have the same genotype, are not excluded.

### Supplementary Figure 3

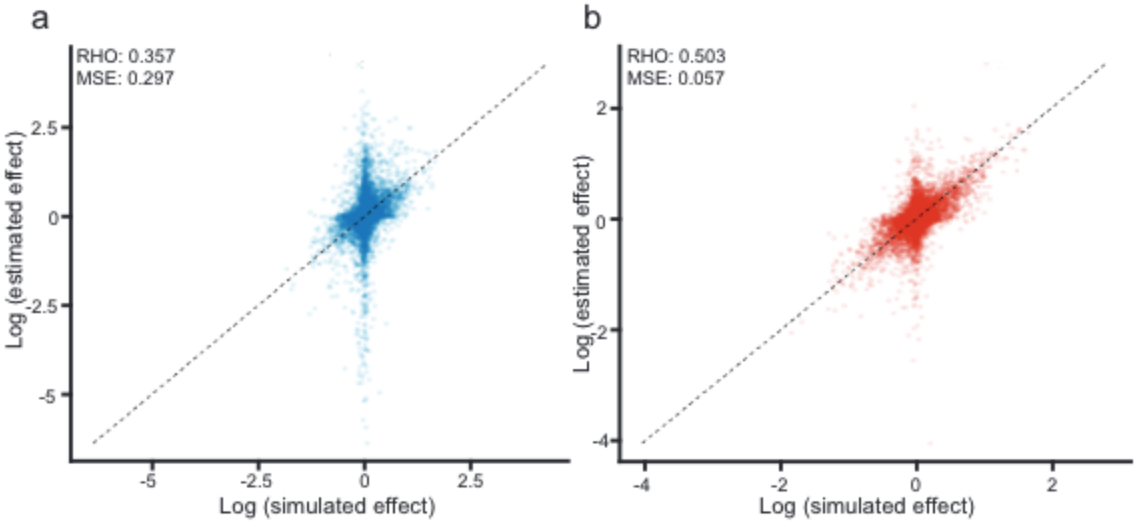


Correlation between simulated effect sizes and effect sizes estimated using the BIRDbath model (a) and the BIRD model (b) , represented in blue and red respectively.  The x-axis represents the log-transformed simulated effect sizes. The y-axis represents the log-transformed estimated effect sizes from the simulated data. The spearman correlation coefficient between these values using the BIRDbath model is 0.503, and the mean squared error of the estimate is 0.057.  The spearman correlation coefficient between these values using the BIRD model is 0.357, and the mean squared error of the estimate is 0.297.

### Supplementary Figure 4

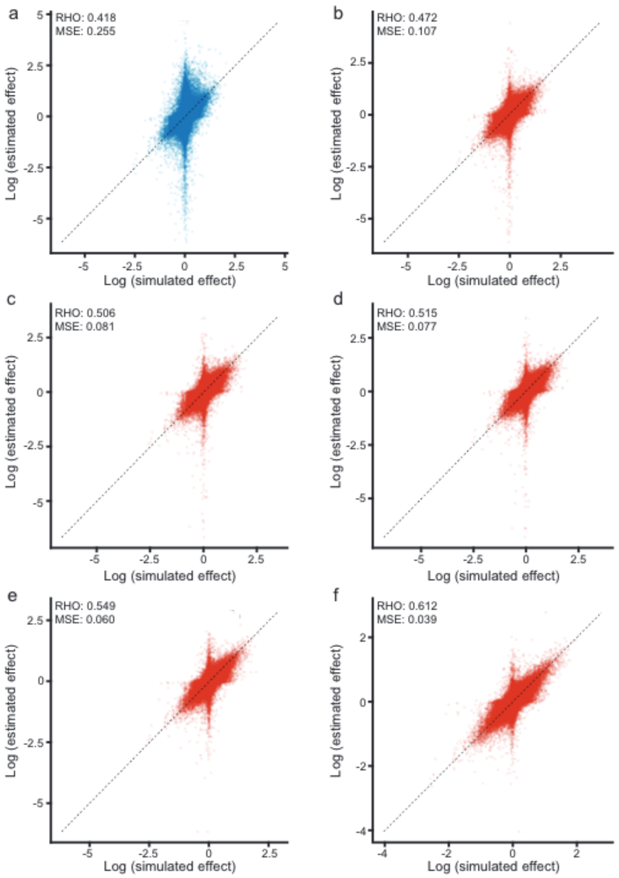


Correlation between simulated effect sizes and effect sizes estimated using the BIRDbath model when varying the number of pools in the simulated experiment. The x-axis represents the log-transformed simulated effect sizes. The y-axis represents the log-transformed estimated effect sizes from the simulated data. In all simulations, there are 100 total samples across all pools. These are divided into (a) 1 pool of 100, (b) 2 pools of 50, (c) 4 pools of 25, (d) 5 pools of 20, (e) 10 pools of 10, and (f) 20 pools of 5. The accuracy of estimates is assessed using the spearman correlation coefficient between the estimated and simulated effects, and the mean squared error, reported for each pooling structure on the corresponding panel.

### Supplementary Figure 5

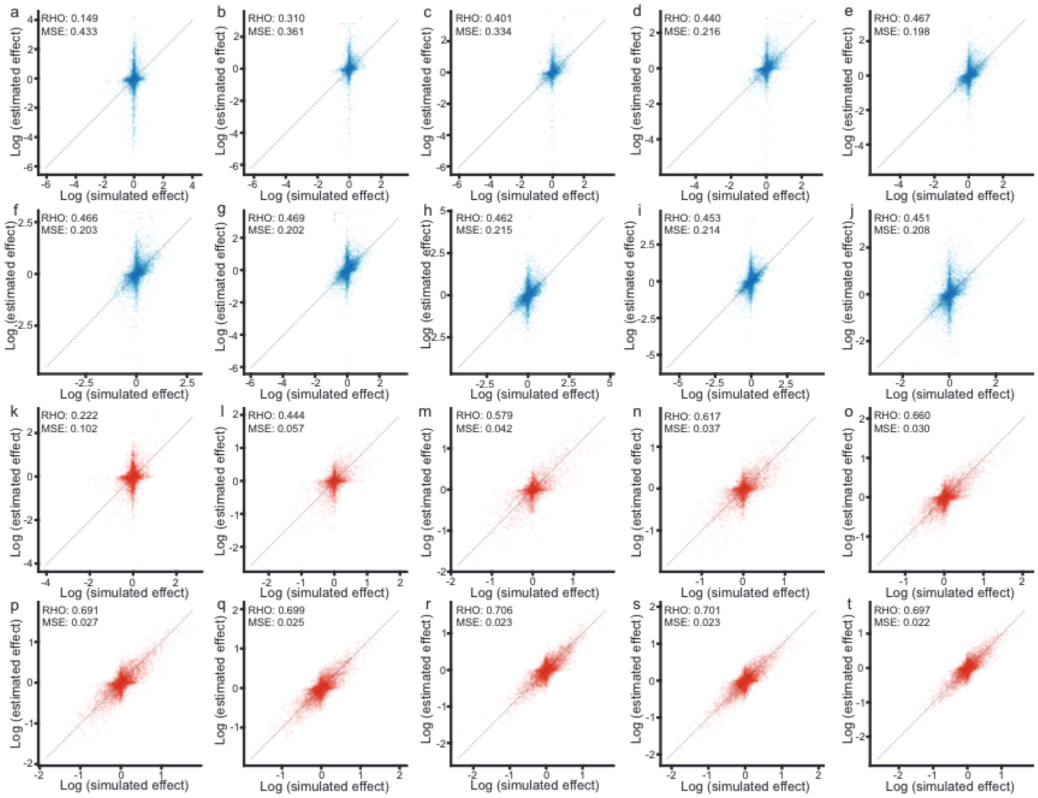

Correlation between simulated effect sizes and effect sizes estimated using the BIRDbath model when varying the number of pools in the simulated experiment. The x-axis represents the log-transformed simulated effect sizes. The y-axis represents the log-transformed estimated effect sizes from the simulated data. In all simulations, there are 100 total samples across all pools. These are divided into (a) 1 pool of 100, (b) 2 pools of 50, (c) 4 pools of 25, (d) 5 pools of 20, (e) 10 pools of 10, and (f) 20 pools of 5. The accuracy of estimates is assessed using the spearman correlation coefficient between the estimated and simulated effects, and the mean squared error, reported for each pooling structure on the corresponding panel.

### Supplementary Figure 6

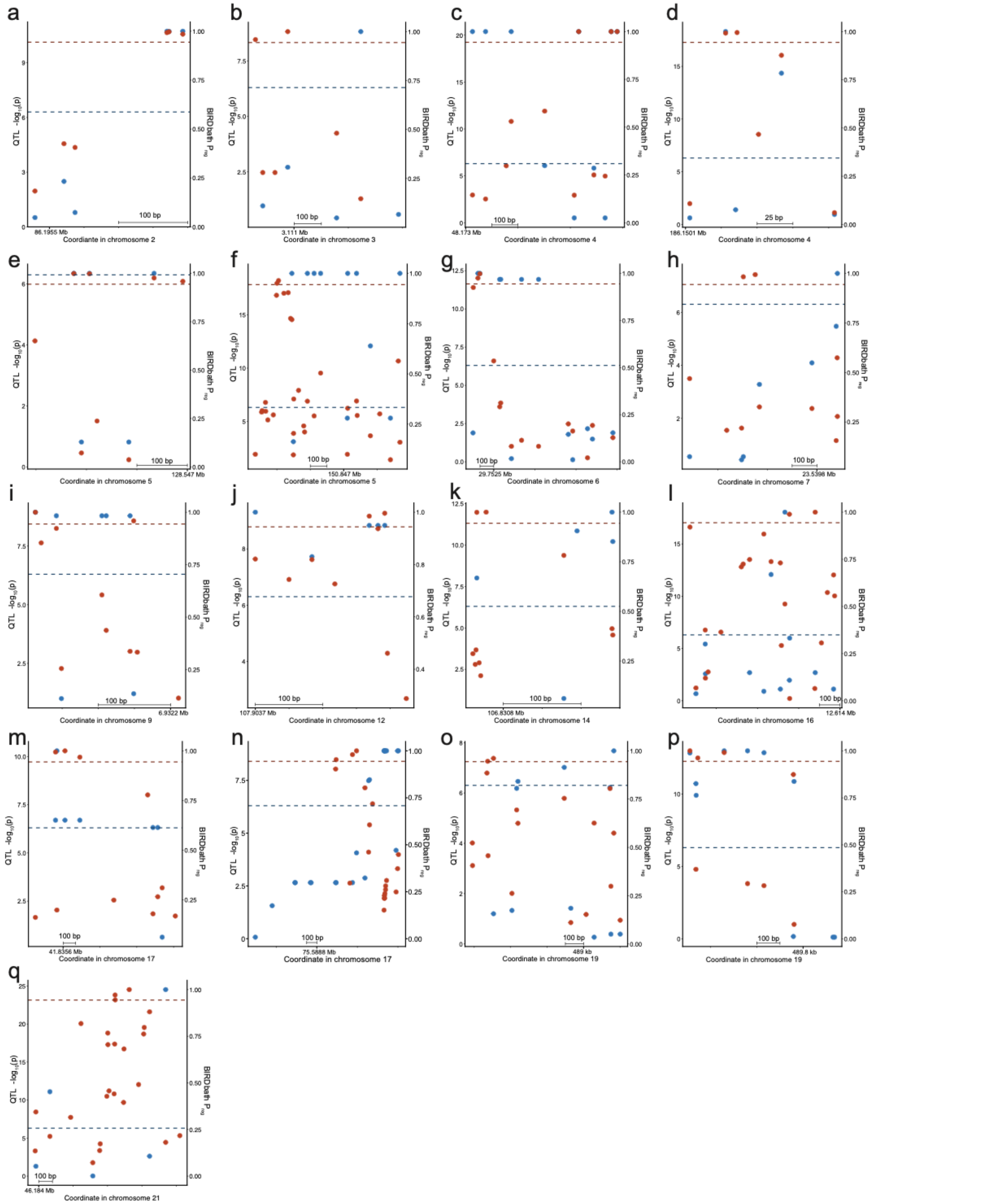

Functional annotation figures comparing the resolution of significant QTLs and STARR-seq variants. The X-axis represents the chromosomal coordinate. Blue points are QTL p-values, represented on the left Y-axis. Red points are STARR-seq $P_{reg}$ values, represented on the right Y-axis. All regions depicted show annotations of caQTLs with multiple significant STARR-seq variants.

#### Supplementary Figure 7

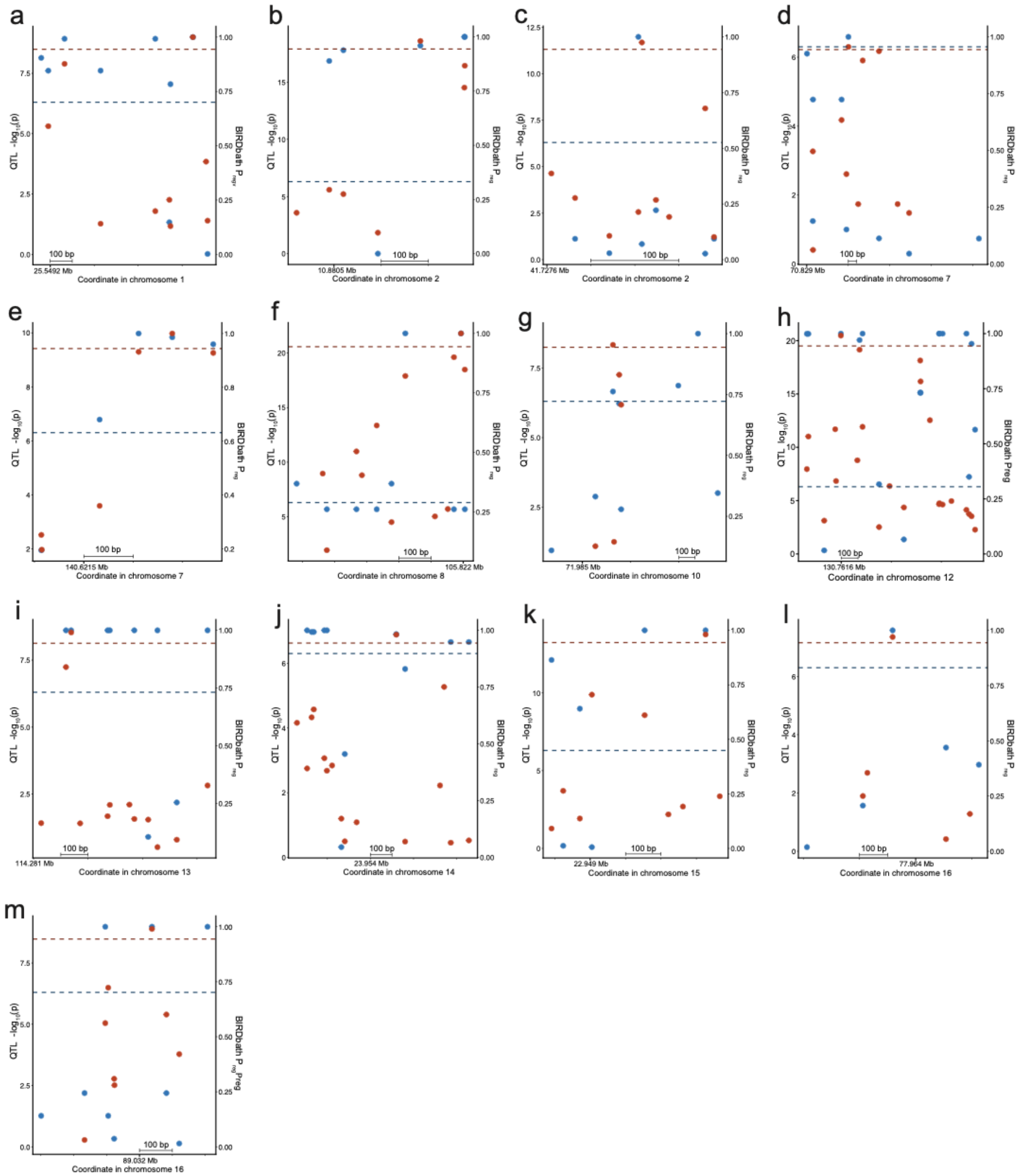


Functional annotation figures comparing the resolution of significant QTLs and STARR-seq variants. The X-axis represents the chromosomal coordinate. Blue points are QTL p-values, represented on the left Y-axis. Red points are STARR-seq $P_{reg}$ values, represented on the right Y-axis. All regions depicted show annotations of caQTLs with significant STARR-seq variants which overlaps a significant caQTL.

### Supplementary Figure 8

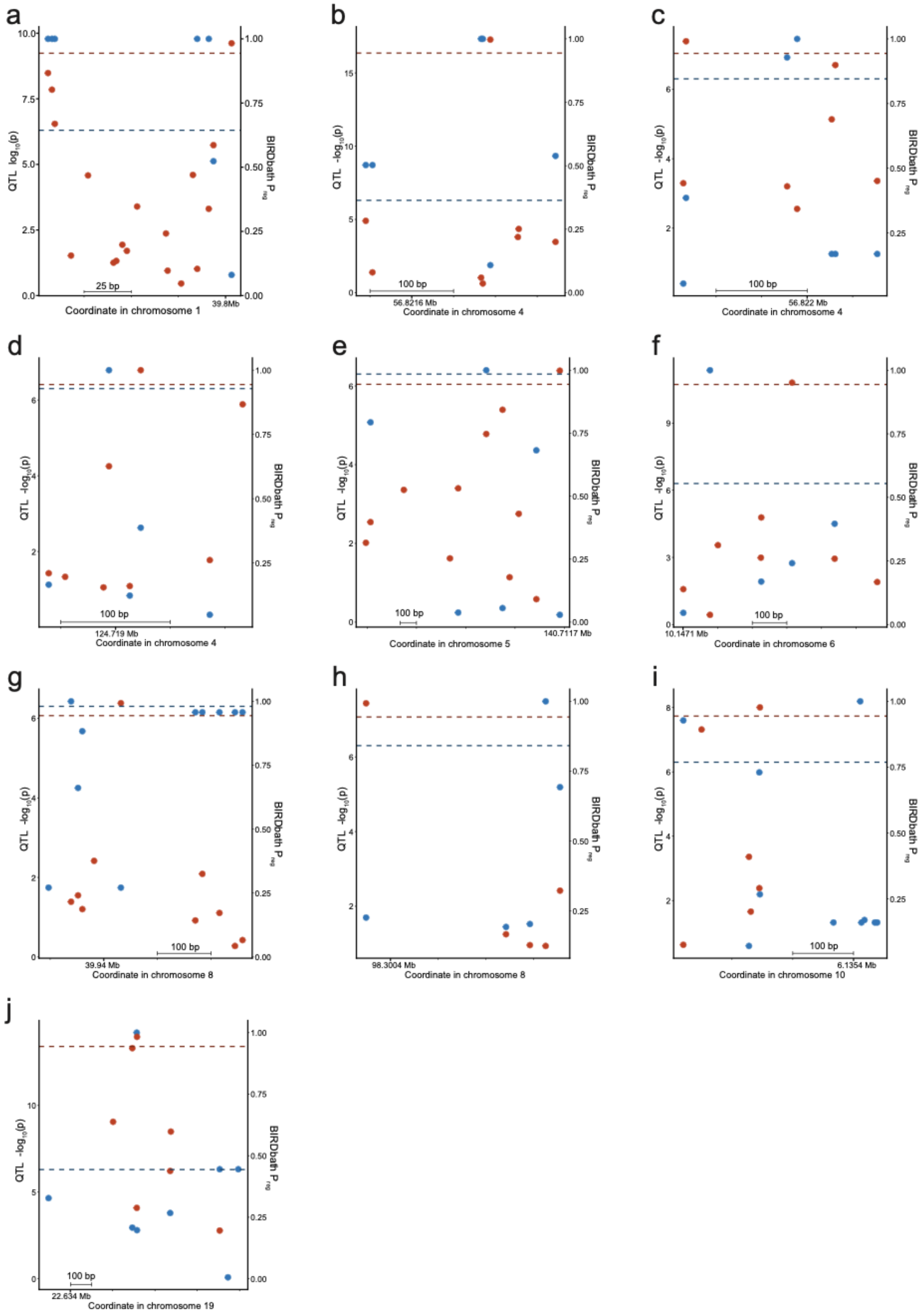


caQTL non- Functional annotation figures comparing the resolution of significant QTLs and STARR-seq variants. The X-axis represents the chromosomal coordinate. Blue points are QTL p-values, represented on the left Y-axis. Red points are STARR-seq $P_{reg}$ values, represented on the right Y-axis. All regions depicted show annotations of caQTLs with a significant STARR-seq variant which overlaps a non-significant caQTL.

### Supplementary Figure 9

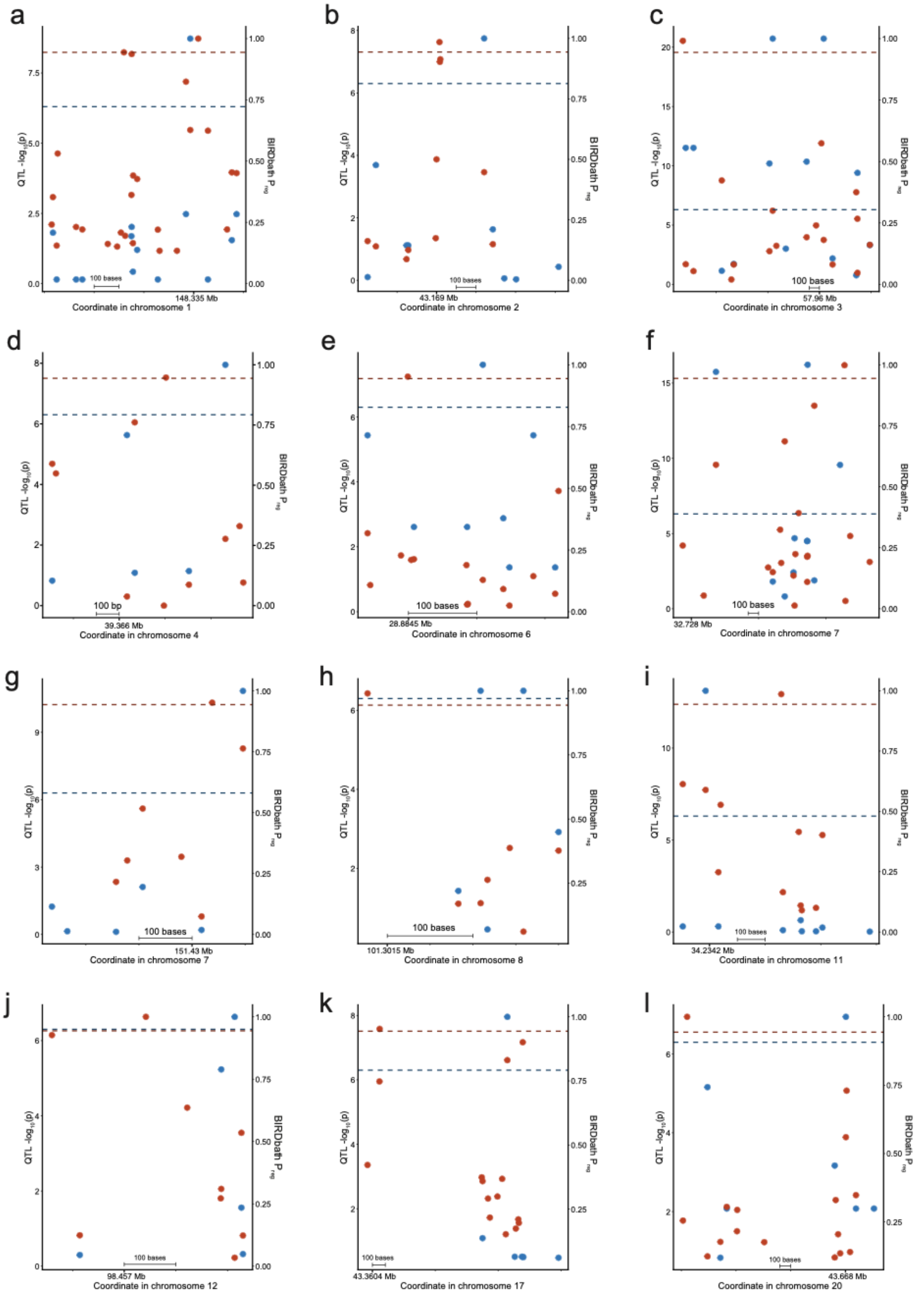

Functional annotation figures comparing the resolution of significant QTLs and STARR-seq variants. The X-axis represents the chromosomal coordinate. Blue points are QTL p-values, represented on the left Y-axis. Red points are STARR-seq $P_{reg}$ values, represented on the right Y-axis. All regions depicted show annotations of caQTLs with a significant STARR-seq variant which does not overlap a caQTL.

### Supplementary Figure 10

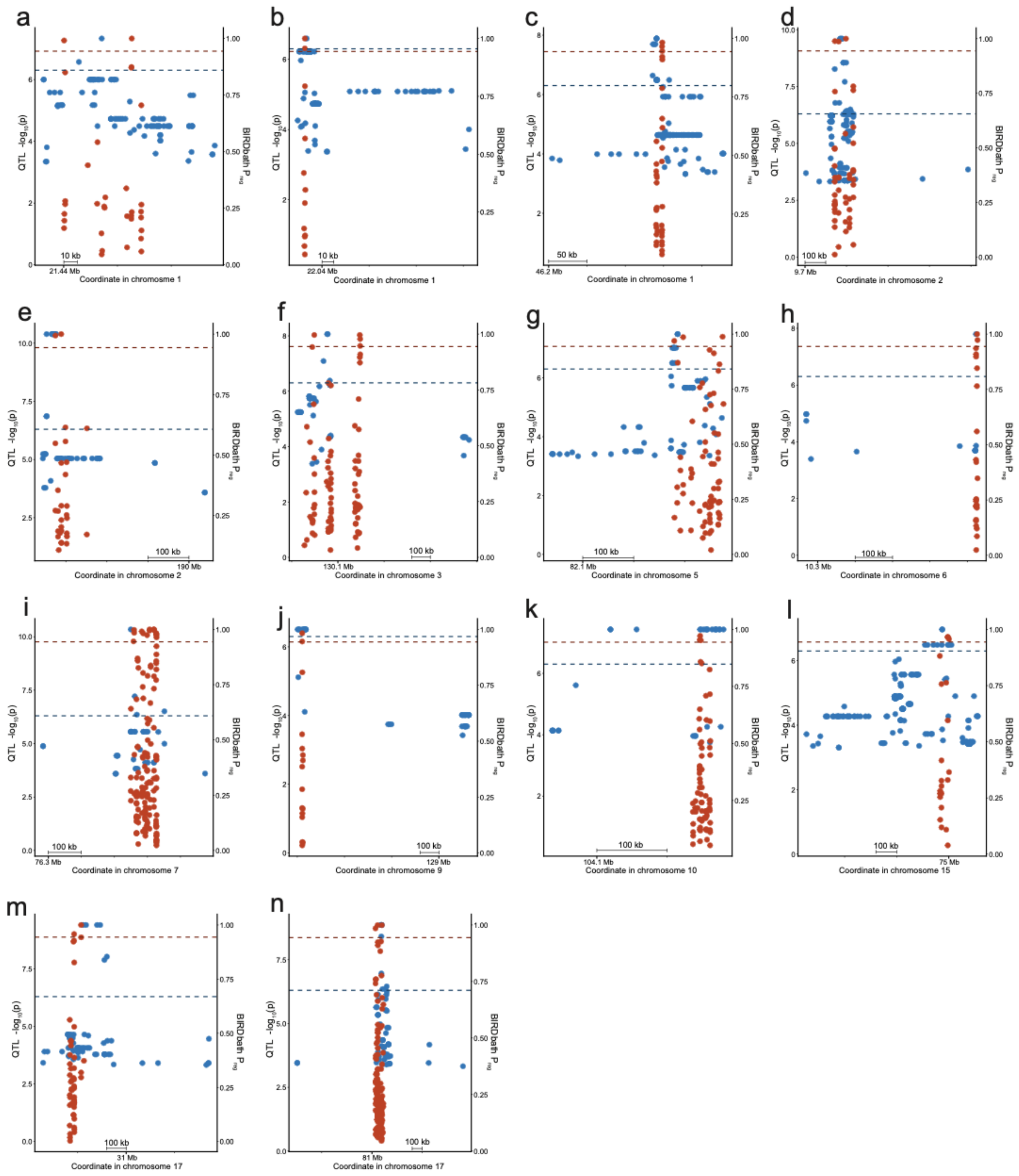


Functional annotation figures comparing the resolution of significant QTLs and STARR-seq variants. The X-axis represents the chromosomal coordinate. Blue points are QTL p-values, represented on the left Y-axis. Red points are STARR-seq $P_{reg}$ values, represented on the right Y-axis. All regions depicted show annotations of eQTLs with multiple significant STARR-seq variants.

### Supplementary Figure 11

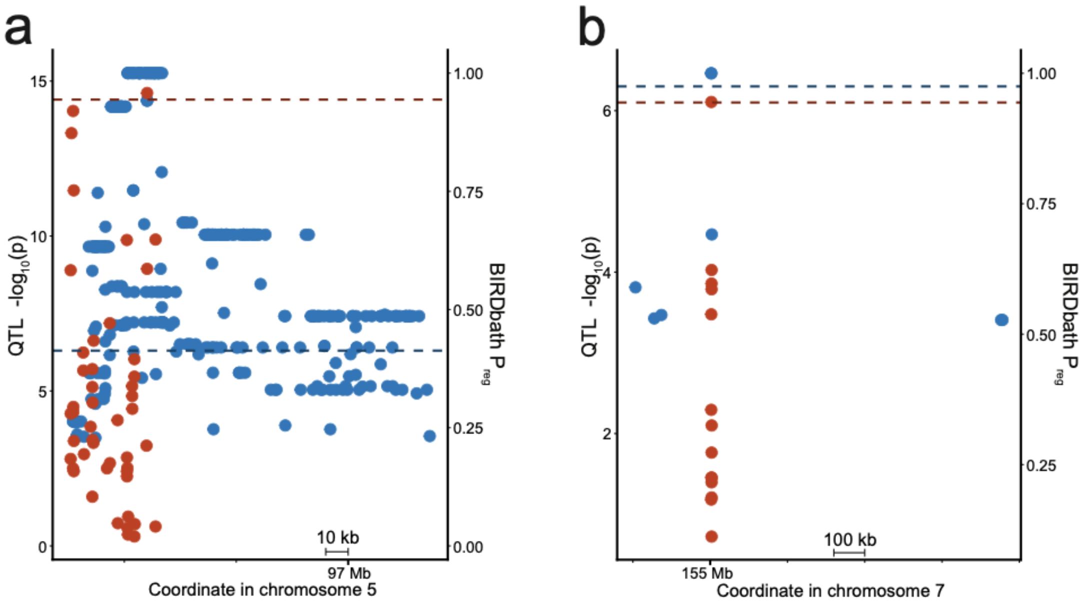


Functional annotation figures comparing the resolution of significant QTLs and STARR-seq variants. The X-axis represents the chromosomal coordinate. Blue points are QTL p-values, represented on the left Y-axis. Red points are STARR-seq $P_{reg}$ values, represented on the right Y-axis. All regions depicted show annotations of eQTLs with a significant STARR-seq variant which overlaps a significant eQTL.

### Supplementary Figure 12

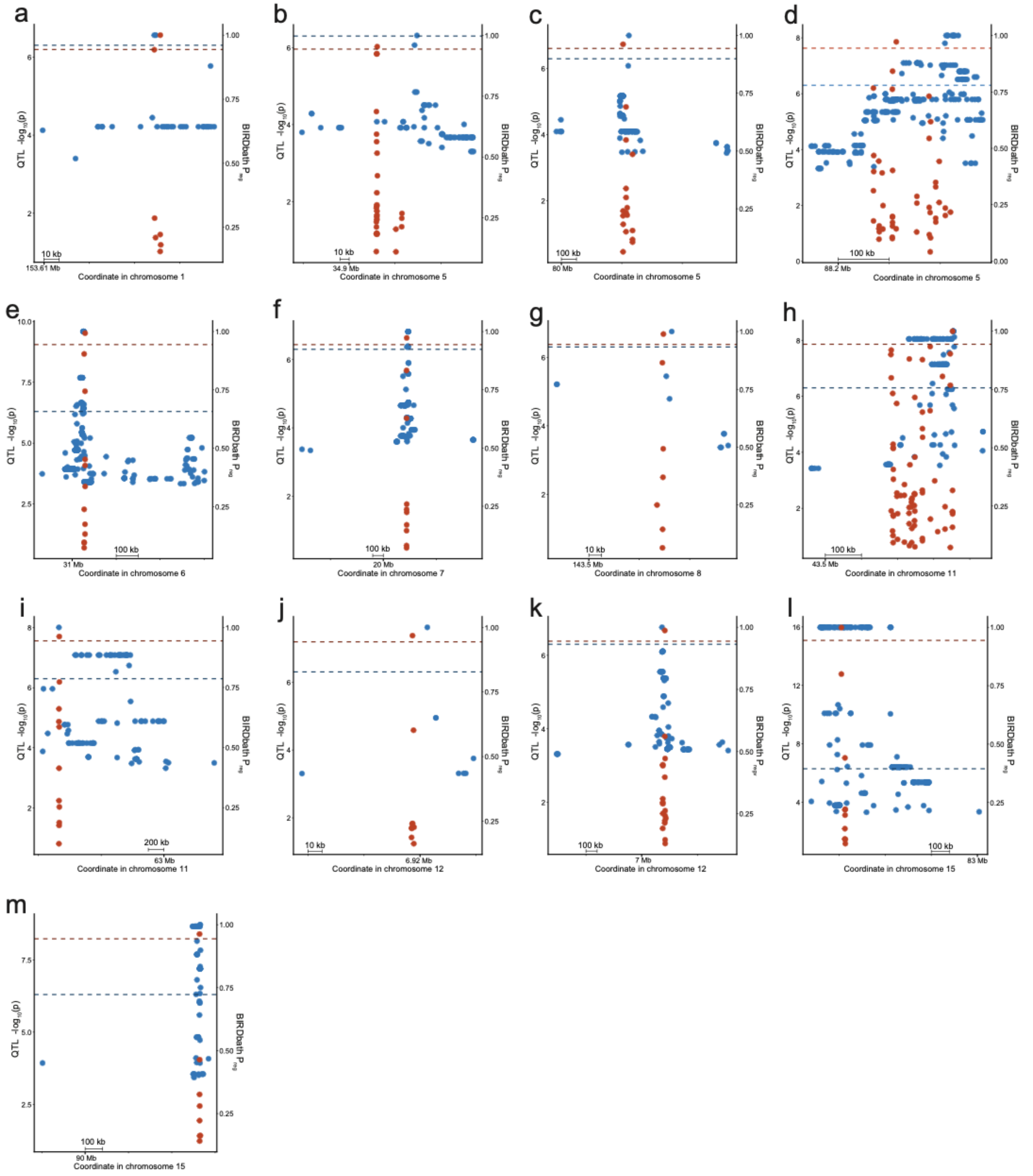


Functional annotation figures comparing the resolution of significant QTLs and STARR-seq variants. The X-axis represents the chromosomal coordinate. Blue points are QTL p-values, represented on the left Y-axis. Red points are STARR-seq $P_{reg}$ values, represented on the right Y-axis. All regions depicted show annotations of eQTLs with a significant STARR-seq variant which does not overlap an eQTL.

### Supplementary Text

#### S1. A realistic simulator for STARR-seq data

We created a simulator for STARR-seq to test the BIRDbath model. It functions by sampling variables in the order that they might be created during an actual experiment. Wherever possible, this dataset was generated from empirical distributions given samples from the Thousand Genomes Project, and distributions observed from pool 1. The counts of each variant are simulated independently for all variants present in pool 1.

First, samples that make up each pool are selected to determine the genotype frequencies of the variants. For the simulations reported in this manuscript, we used the 100 samples which were present in the multi-pool experiment. For each variant, the genotype within the pools was sampled jointly with the expected effect size of the variant given the results of running the BIRD model on the counts generated by pool 1.

The measured DNA frequency was sampled from the distribution of measured allele frequencies in pool 1, conditional on the expected allele frequency in the pool. The expected allele frequency in the RNA was calculated based on the measured allele frequency in the DNA and the expected effect size:

$$q = \frac{\theta \times p}{1 - p + (\theta \times p)}$$

where $\theta$ is the effect size of the variant and $p$ is the measured allele frequency in the DNA. The measured allele frequency in the RNA is sampled from a beta distribution

$$q_{beta} \sim beta(\mathrm{mode} = q, \mathrm{beta} = \mathrm{conc})$$

where $q_{beta}$ is the measured allele frequency in the RNA, $q$ is the previously calculated value for the expected allele frequency in the RNA, and $\mathrm{conc}$ is a concentration parameter that was previously calculated[23]. Here, we use 139 for the value of that concentration parameter.

The total coverage of the DNA library was sampled from the known distribution of genomic coverages in pool 1. The ratio of the RNA coverage to DNA coverage was calculated separately for both the reference and alternate alleles using known data from pool 1:

$$R_a = k_o / a_o$$

$$R_r = m_o / b_o$$

where $a_o$ and $k_o$ are previously observed alternate allele counts in the DNA and RNA respectively, and $b_o$ and $m_o$ are previously observed reference allele counts in the DNA and RNA respectively. From those two ratios, a single overall coverage ratio was calculated as the weighted average of those two values with the weighting given by the genotype of the simulated pool:

$$R = R_a(g) + R_r(1 - g)$$

where $R_a$ and $R_r$ are the previously calculated alternate and reference allele ratios, and $g$ is the expected frequency of the alternate allele in the DNA. This overall ratio was used with the simulated total DNA coverage to calculate the simulated total RNA coverage:

$$RNA_c = DNA_c \times R$$

where $RNA_c$ and $DNA_c$ are the coverage values in the RNA and DNA respectively, and $R$ is the weighted expected ratio of the two libraries.

The alternate allele counts are generated from a binomial distribution, and the reference allele counts are calculated as the difference between the alternate allele count and the total coverage. For the DNA libraries, this is sampled as follows:

$$a \sim \text{binomial}(DNA_c, p)$$
$$b = DNA_c - a$$

where $DNA_c$ is the sampled coverage in the DNA library, $p$ is the measured alternate allele frequency, and $a$ and $b$ are the simulated counts in the DNA library for the alternate and reference alleles respectively. Similarly, the allele counts in the RNA libraries are sampled as follows:

$$k \sim \text{binomial}(RNA_c, q_{beta})$$
$$m = RNA_c - k$$

Where $RNA_c$ is the calculated coverage in the RNA library, $q_{beta}$ is the measured alternate allele frequency, and $k$ and $m$ are the simulated counts in the RNA library for the alternate and reference alleles respectively.

In the case where the expected allele frequency in the pool is 0, the measured alternate allele frequency in both the DNA and RNA libraries are also 0. The alternate allele count in both libraries is set to 0, and the reference allele count in both libraries is equal to the coverage. Similarly, when the expected alternate allele frequency is 1, the alternate allele counts are equal to the coverage, and the reference allele counts are equal to 0.

This simulation was repeated for every variant present in pool 1.

### S2. A simulator to examine how heterogeneity changes with STARR-seq experimental design

The previous simulator was modified to assess how the heterogeneity in the experiments compares between a multi-pool experiment and many replicates of the same experiment. The simulator to answer this question is very similar to the simulator described in Supplemental Text S1, with a few modifications.

The samples used to determine pool genotypes were previously based on the samples used in the multi-pool experiments, but here they are selected randomly.

The number of pools was kept constant in the previous simulation, but here it changes to show how the heterogeneity can vary. Because the total number of samples is held constant at 100 genotypes in both simulations, this change to the number of pools also changes the size of the pools, and the assignment of the samples to pools. This simulation is repeated for any number of pools from 1, where all 100 samples would be in the same pool, to 100, where each sample would be in its own pool. The samples were randomly assigned to pools so that the multi-pool simulations with pools that divided unevenly had a difference of at most one individual between any two pools.

Because these were no longer based entirely on empirical distributions, the coverage in the DNA and RNA libraries was no longer sampled from the empirical distributions. In this simulation, the DNA coverage in each DNA replicate and in each RNA replicate are equal to 1000 reads.

Finally, this simulation was created to compare the heterogeneity in a multi-pool experiment and in a single pool experiment, so it has to simulate both multi-pool and single-pool experiments which are comparable.

The DNA and RNA libraries for multi-pool experiments were simulated like in the simulator described in Supplementary Text S1, where each pool had a different DNA library simulated, and the RNA library was based on that DNA library.

When a single-pool experiment was being simulated, only one DNA library was generated by the simulator for any n number of replicates. The expected alternate allele frequency for this simulation was randomly selected from n different pool options. The counts for this DNA library were simulated only once. The RNA libraries were simulated n times, all based on the measured alternate allele frequency of that one DNA library.

When running this simulation, one variant is chosen to determine all genotypes. The effect size was set to $\theta = 0.5$. This simulation was repeated by dividing $i$ times for $i \in \{1, 100\}$, and each division was simulated 100 times.

The heterogeneity of these samples was compared, defining heterogeneity of a sample as the sample variance among the RNA frequencies of a single multi-pool or single-pool experiment. The heterogeneity of each repetition of each simulated split is reported, along with the average heterogeneity value for all repetitions of a single set of splits.

### S3. A simulator to demonstrate the effect of heterogeneity on effect estimation

Finally, we tested the effect of the heterogeneity on effect estimation. Here, the simulator was modified from the simulator described in Supplementary Text S2.

100 random genotypes from the Thousand Genomes Project were selected again, but now they were only split into up to 10 pools.

The total coverage in the DNA and in the RNA were held at 1000 reads, and the exact coverage of each replicate was variable. These coverage values were calculated as in Majoros et.al. 2020 (BIRD PMID), where the smallest replicate was not less than 62% of the largest replicate. Specifically:

$$M_1 = 0.62\, M_R$$
$$M_i = M_1 + i\beta$$
$$\sum_{i=1}^{R} M_i = N_{RNA}$$

where $M_1$ is the coverage in the smallest replicate, $M_R$ is the largest replicate, $i$ is the number of the replicate, $R$ is the total number of replicates, $\beta$ is a slope value, and $N_{RNA}$ is the total RNA value.

Four metrics are reported to assess the impact of heterogeneity. The variance in the total RNA alternate allele count and alternate allele frequency are defined as the sample variance of the total RNA alternate allele count or frequency across all splits of a single simulation. The effect size was calculated using the naive estimate:

$$\theta_{naive} = \frac{k/a}{m/b}$$

where $\theta_{naive}$ is the naive effect size, $k$ and $m$ are the counts in the RNA of the alternate and reference alleles respectively, and $a$ and $b$ are the counts in the DNA library of the alternate and reference alleles respectively. The sample variance of this value across all simulations is reported. Finally, the RMSE of the estimate is defined as the root mean squared error of the estimate relative to the simulated value.

This simulation was repeated for 10,000 simulated variants. When splitting the pools, sometimes the one selected for the replicate simulation had an alternate allele frequency that was very different from the simulated overall alternate allele frequency. To mitigate the effect that this sampling would have on the simulation, these simulations were repeated 30 times, each time with different splits of the genotypes, and thus different alternate allele frequencies in the individual pools or replicates.
